# The role of a population of descending neurons in the optomotor response in flying *Drosophila*

**DOI:** 10.1101/2022.12.05.519224

**Authors:** Emily H. Palmer, Jaison J. Omoto, Michael H. Dickinson

## Abstract

To maintain stable flight, animals continuously perform trimming adjustments to compensate for internal and external perturbations. Whereas animals use many different sensory modalities to detect such perturbations, insects rely extensively on optic flow to modify their motor output and remain on course. We studied this behavior in the fruit fly, *Drosophila melanogaster*, by exploiting the optomotor response, a robust reflex in which an animal steers so as to minimize the magnitude of rotatory optic flow it perceives. Whereas the behavioral and algorithmic structure of the optomotor response has been studied in great detail, its neural implementation is not well-understood. In this paper, we present findings implicating a group of nearly homomorphic descending neurons, the DNg02s, as a core component for the optomotor response in flying *Drosophila*. Prior work on these cells suggested that they regulate the mechanical power to the flight system, presumably via connections to asynchronous flight motor neurons in the ventral nerve cord. When we chronically inactivated these cells, we observed that the magnitude of the optomotor response was diminished in proportion to the number of cells silenced, suggesting that the cells also regulate bilaterally asymmetric steering responses via population coding. During an optomotor response, flies coordinate changes in wing motion with movements of their head, abdomen, and hind legs, which are also diminished when the DNg02 cells are silenced. Using two-photon functional imaging, we show that the DNg02 cells respond most strongly to patterns of horizontal motion and that neuronal activity is closely correlated to motor output. However, unilateral optogenetic activation of DNg02 neurons does not elicit the asymmetric changes in wing motion characteristic of the optomotor response to a visual stimulus, but rather generates bilaterally symmetric increases in wingbeat amplitude. We interpret our experiments to suggest that flight maneuvers in flies require a more nuanced coordination of power muscles and steering muscles than previously appreciated, and that the physical flight apparatus of a fly might permit mechanical power to be distributed differentially between the two wings. Thus, whereas our experiments identify the DNg02 cells as a critical component of the optomotor reflex, our results suggest that other classes of descending cells targeting the steering muscle motor neurons are also required for the behavior.

## INTRODUCTION

As flying animals navigate through space, they must constantly integrate sensory information to ensure they maintain stable flight along their desired trajectory. Studies on the behavioral algorithms that flying insects use to adjust for internal and external perturbations have demonstrated their ability to maintain constant groundspeed in the face of variable wind speed (David, 1978; Kennedy, 1940; Leitch et al., 2021), reject applied body rotations (Beatus et al., 2015; Ristroph et al., 2010; Whitehead et al., 2015), and sustain flight despite having extensive wing damage (Muijres et al., 2017). Much recent research on flight control has focused on the fruit fly *Drosophila melanogaster* due to its robust behavior in tethered flight arenas (Götz, 1987) and the vast array of genetic tools available in this species (Simpson and Looger, 2018). Collectively, these efforts provide a broad overview on the basic algorithmic structures of insect flight control systems (Dickinson and Muijres, 2016).

To study flight stabilization, researchers often exploit the optomotor response, a well-characterized behavior in which a fly reflexively steers so as to minimize the optic flow it perceives. The optomotor response is typically quantified by measuring either the yaw torque generated by the wings (Götz, 1968) or the bilateral difference of the wingbeat amplitude (Götz et al., 1979), which is strongly correlated with torque (Tammero et al., 2004). The motor output induced by patterns of wide-field optic flow has been extensively studied, including a detailed description of the responses in flight muscles. The flight muscles of flies are functionally stratified into two systems that collectively power and regulate wing motion (Dickinson, 2006). The first system consists of large asynchronous muscles that provide the power to flap the wings at high frequency. These stretch-activated muscles (Josephson et al., 2000; Pringle, 1949) insert on the walls of the thorax in an approximately orthogonal arrangement, creating a self-oscillatory system that drives the back and forth motion of the wings. Dorsal longitudinal muscles (DLMs) contract to move the wing forward while stretching and activating dorso-ventral muscles (DVMs), which subsequently drive the wings backward while stretching the DLMs to continue the cycle. The oscillations of the power muscles are transmitted to the wings via their action on two critical mechanical inputs to the wing hinge: the scutellar lever arm and the anterior notal wing process (Boettiger and Furshpan, 1952; Pringle 1957; Miyan and Ewing, 1984; Deora et al., 2015). The second system consists of tiny synchronous steering muscles that actuate subtle changes in wing kinematics by directly reconfiguring the wing hinge (Dickinson and Tu, 1997; Heide, 1968; Heide and Götz, 1996; Lindsay et al., 2017). Whereas this prior work has highlighted the important role of steering muscles in generating the asymmetries in wing motion necessary for the optomotor response, the power muscles are generally thought to operate in a bilaterally symmetric fashion because the scutellar lever arms are mechanically coupled across the two sides of the fly (Boettiger and Furshpan, 1952; Deora et al., 2015; Pringle, 1957; Walker et al., 2014). However, bilaterally asymmetric Ca^2+^ signals recorded from the power muscles during presentation of visual yaw stimuli suggest that they, too, might play a role in the optomotor response (Lehmann et al., 2013). Thus, the changes in wing motion during optomotor reflexes could involve a coordination of both power and steering muscle activity. Such coordination is not unexpected, given that the power requirements to flap a wing depend on the torque that it generates (Ellington, 1984), which must change during the optomotor response. The question remains, however, whether there is enough flexibility in the mechanical structures linking the scutellar level arms and anterior notal wing processes on the two sides of the fly such that asymmetric activity in the power muscles could generate differential drive to the left and right wing hinges.

Whereas the optomotor response has been studied in great detail at the behavioral level, the details of the sensory-motor pathway that underlie it remain largely unknown, especially the means by which wide-field visual motion is conducted to the motor centers in the ventral nerve cord (VNC). Recent work morphologically classifying a large fraction of the descending neurons (DNs) provides a useful starting point for investigating how sensory information from the brain is transformed into motor commands in the VNC (Namiki et al., 2018). Namiki and colleagues (Namiki et al., 2022) recently performed an optogenetic activation screen on a set of driver lines selectively targeting different DNs and identified a class of nearly homomorphic neurons, the DNg02s, as a candidate for encoding commands relevant to flight control. Unlike many identified DNs, which exist as single bilateral pairs of the cells with stereotyped morphology, the DNg02 cells constitute a large set of nearly homomorphic cells. The potential role of the DNg02 cells in flight control was identified via optogenetic activation of the neurons in tethered flying animals, using a collection of driver lines that targets different numbers of DNg02 cells. The change in bilateral stroke amplitude elicited by the optogenetic stimulus was linearly correlated with the number of cells activated, suggesting that the fly might employ the DNg02s in a population code to regulate wingbeat amplitude and total mechanical power over a large operating range. One hypothesis for the large number of DNg02 cells is that they collectively provide both the dynamic range and the motor precision that is required for flight control. Whereas the ability to fly straight with damaged wings requires that flies maintain large bilateral differences in wing motion (Muijres et al., 2017), these kinematics must be regulated with extreme precision because of the strong nonlinear influence of wing length and speed on aerodynamic forces and moments (Ellington, 1984).

In this paper, we conducted extensive experiments to further elucidate the role of the DNg02 neurons in flight stabilization. By genetically silencing the DNg02 cells, we show that the magnitude of the optomotor response is diminished in proportion to the number of pairs silenced, providing support for the population coding hypothesis. These results indicate that these neurons are necessary for the optomotor response, insofar as they are required for the response to reach its full magnitude. We employed two-photon functional imaging of DNg02 neurons in an array of different driver lines, confirming that this cell class exhibits bilaterally asymmetric responses to visual yaw stimuli (Namiki et al., 2022). However, when we performed unilateral activation of the neurons, we found that this manipulation does not elicit turning responses; rather, we observed bilaterally symmetric increases in wingbeat amplitude upon unilateral activation. This result suggests that additional pathways are likely involved in the optomotor response, perhaps involving DNs that make strong connections to steering muscle motor neurons. Our results thus support the hypothesis that the optomotor response involves a complex coordination of both power and steering muscle activity, underscoring the sophisticated nature of the insect wing hinge and its control.

## RESULTS

### Coordination of different motor systems during the optomotor response

To probe the behavioral structure and neuronal implementation of the optomotor response, we tracked the kinematics of tethered flies while presenting open-loop visual stimuli simulating rotation about the yaw axis (Figure 1A-B). We aligned flying flies to a machine vision system that measured wingbeat amplitude in real time, whereas head angle, abdomen angle, and leg positions were measured offline from the recorded image stream using DeepLabCut (Mathis et al., 2018). An example of a behavioral response to wide field visual motion is plotted in Figure 1C. The optomotor response is often quantified in tethered flies using the bilateral difference in wingbeat amplitude of the two wings, ΔWBA. Under the assumption that the flies’ behavior is bilaterally symmetric, we normalized these responses such that the angle of the wing on the inside of the induced turn (WBA_i_) was subtracted from the angle of the outside wing (WBA_o_) (Figure 1D), thereby combining yaw responses to the left and right into one data set. Upon presentation of the optomotor stimulus, ΔWBA rapidly increased until it reached a plateau of approximately 20° that was maintained for the duration of the stimulus period. The flies generated this difference in wingbeat amplitude via a decrease in WBA_i_ of ~7°, and an increase in WBA_o_ of ~13° (Figure S1A-B). At the offset of the stimulus, ΔWBA decayed slowly as has been noted previously (Schnell et al., 2014).

**Figure 1.**
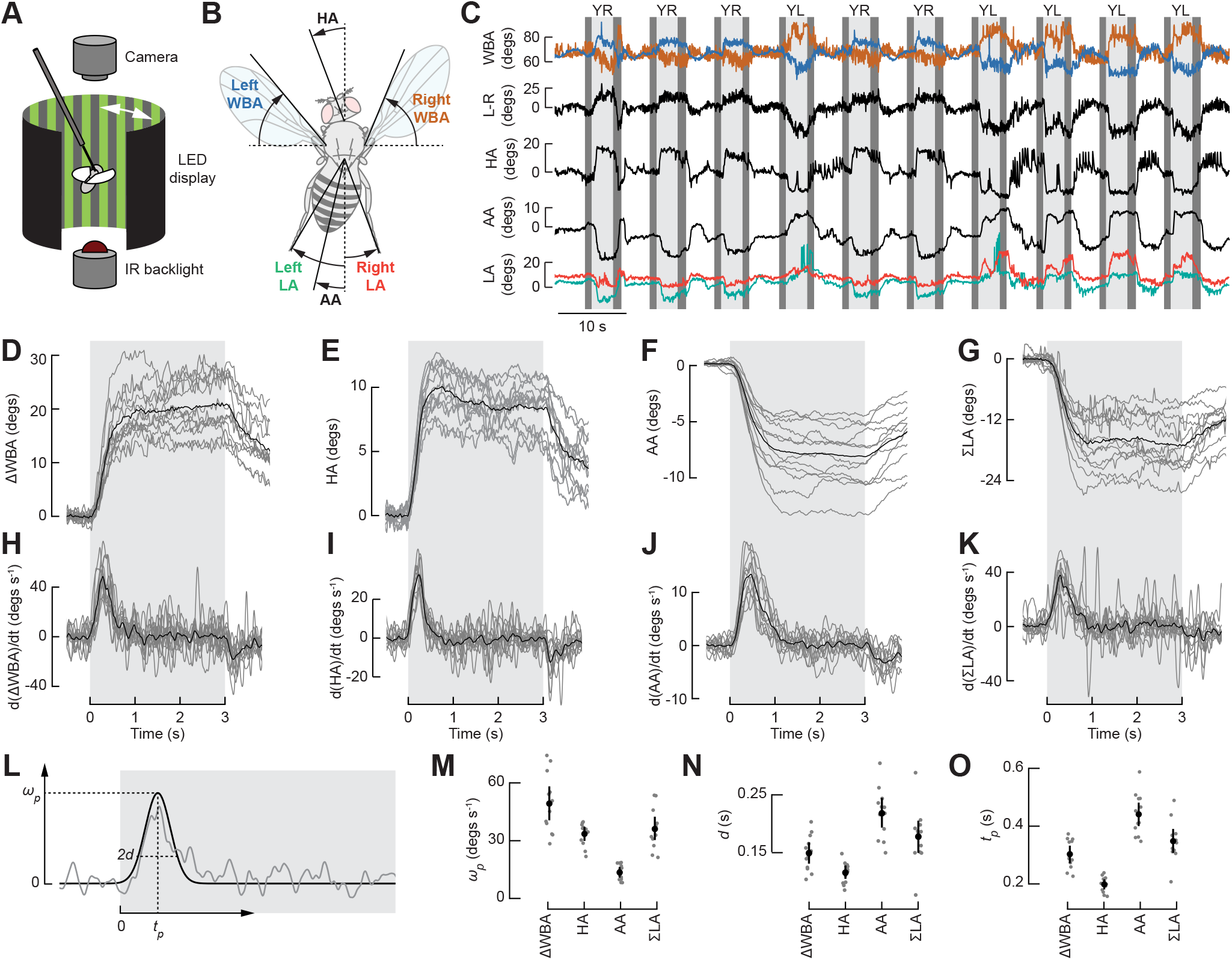
The optomotor response consists of coordinated actions of different motor systems. (A) Schematic of the experimental set-up (not drawn to scale). The fly is centered within a curved visual display of green LEDs. An image of the fly is captured on a downward facing camera and analyzed using a real-time machine vision system that measures the wingbeat amplitude of the left and right wing. (B) Cartoon showing the kinematic parameters recorded during experiments. (C) Example time traces of left and right wingbeat amplitudes (WBA), left minus right amplitude (L-R), head angle (HA), abdomen angle (AA), and leg angles (LA). Presentations of yaw motion to the left (YL) and right (YR) (light gray) were interspersed with epochs of closed loop stripe fixation (white) and a static striped drum pattern (dark gray). The left leg is shift vertically by 20° for visual clarity. (D) Averaged wingbeat amplitude differential (ΔWBA) during the presentation of yaw motion (n = 12 flies). ΔWBA is normalized to the direction of the yaw motion, such that the amplitude of the wing decreasing in amplitude (the inside wing relative to the direction of motion, WBA_i_) is subtracted from the amplitude of the wing increasing in amplitude (the outside wing, WBA_o_). The mean response of individual flies is shown in light gray and the mean across all flies is shown in black. The period of stimulus presentation is shown by the gray patch. (E) As in Figure 1D, for HA. (F) As in Figure 1D, for AA. (G) As in Figure 1D, for the sum of the leg angles (ΣLA). (H) Time derivative of ΔWBA, calculated using a Savitzky-Golay filter (n = 12 flies). Plotting conventions as in Figure 1D. (I) As in Figure 1H, for HA. (J) As in Figure 1H, for AA. (K) As in Figure 1H, for Σ LA. (L) Schematic showing the definitions of the parameters governing the Gaussian functions fit to the time derivative traces in Figure 1H-K. See text for details. (M) Peak angular velocity of ΔWBA, head, abdomen, and leg movements. (N) Duration of wing, the head, abdomen, and leg movements. (O) Timing of wing, the head, abdomen, and leg movements.

Whereas the optomotor response is typically quantified by tracking changes in wing motion, flies coordinate their wing responses with adjustments in head position and body posture. The head motion is thought aid in gaze stabilization by minimizing retinal slip (Cruz et al., 2021; Hengstenberg, 1993; Land, 1973), while ruddering of the legs and abdomen are thought to contribute torque (Berthé and Lehmann, 2015; Götz et al., 1979). The head motion we measured consisted of a quick initial yaw rotation to a plateau value of ~10°, followed by a decay at the termination of the optomotor stimulus (Figure 1E). Similarly, the flies quickly deflected their abdomen and hind legs at the onset of visual motion (Figure 1F-G, Figure S1C-D), consistent with prior studies (Berthé and Lehmann, 2015; Zanker, 1988). These deflections of the abdomen and hind legs have been suggested to complement the torque generated by the wings during the optomotor response (Götz et al., 1979), a hypothesis that is supported by our results that the abdomen and legs deflected toward the side of the inside wing of the fictive turn.

To quantify and compare the dynamics of the different motor reflexes, we calculated the time derivatives of the wing, head, abdomen, and leg responses, and then normalized the sign of the responses such that the peak angular velocity is plotted as positive (Figure 1H-K, Figure S1E-H). In the case of the wing responses, we wish to make clear that we measured the velocity of the changes in ΔWBA (which is an angle representing the left-right difference in the ventral extent of the stroke envelopes), not the angular velocity of the individual wings as they flapped back in forth, which is of course much faster. As all four reflexes consisted of a quick initial response followed by a plateau or slow movement, the derivatives all exhibit an initial peak in angular velocity that is well approximated by a Gaussian function, which provides a convenient means for quantitative comparison. A Gaussian fit parametrizes the behavioral response as

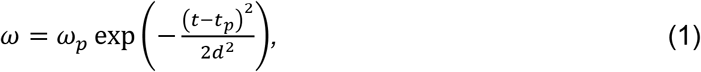

where *ω* is the angular velocity of the given body structure, *t* is the time in seconds, *ω_p_* is the peak angular velocity, *t_p_* is the time of the peak angular velocity, and *d* quantifies the width of the curve (Figure 1L). We found that *ω_p_* was greatest for the ΔWBA and smallest for the abdomen movement (Figure 1M). When the wings and legs are decomposed into the inside and outside appendages individually, we found that the head angle changes with the greatest angular velocity (Figure S1I). Our results are consistent with prior work (Duistermars, 2012; Schilstra and van Hateren, 1998) showing that the duration of the head movement in flies is less than that of the wings (Figure 1N, Figure S1J). The head movement peaks the earliest (Figure 1O, Figure S1K), suggesting that, as with the optomotor response of walking flies (Cruz et al., 2021), head movement precedes that of the rest of the body. The rapidity of head movement relative that we measured in these fictive maneuvers is also consistent with studies of spontaneous saccades in free flying flies (Schilstra and van Hateren, 1998; Wagner and Land, 1986).

### Consistent morphological variation among DNg02 cells suggests the existence of distinct sub-classes

Because DNg02 neurons appear well suited to regulate wing motion over a large dynamic range via a population code and receive input from regions of the brain associated with visual processing (Namiki et al., 2018; Namiki et al., 2022), we sought to examine their potential role in the optomotor response. To this end, we first set out to conduct a more comprehensive anatomical analysis of the 13 available split-Gal4 driver lines, to better interpret the results of experimental manipulations. High resolution confocal imaging of preparations expressing both membrane-bound GFP and nuclear DsRed enabled us to visualize the fine arbors and more reliably count the number of somata in each line (Figure 2A-B, S2A). The cell bodies of DNg02 neurons reside at the ventral edge of the gnathal ganglion (GNG), with axons forming a fiber tract that runs ventrally along the GNG before ascending dorsally (Figure 2B). To be included in our quantification of cell number, a cell body was required to have a nucleus in the appropriate cluster and an axon following the relevant tract. On average, we found 40% more DNg02 cells in each driver line than reported previously (Namiki et al., 2018; Namiki et al., 2022), a discrepancy that we attribute primarily to the benefit of marking nuclei with DsRed, which make somata easier to count.

**Figure 2.**
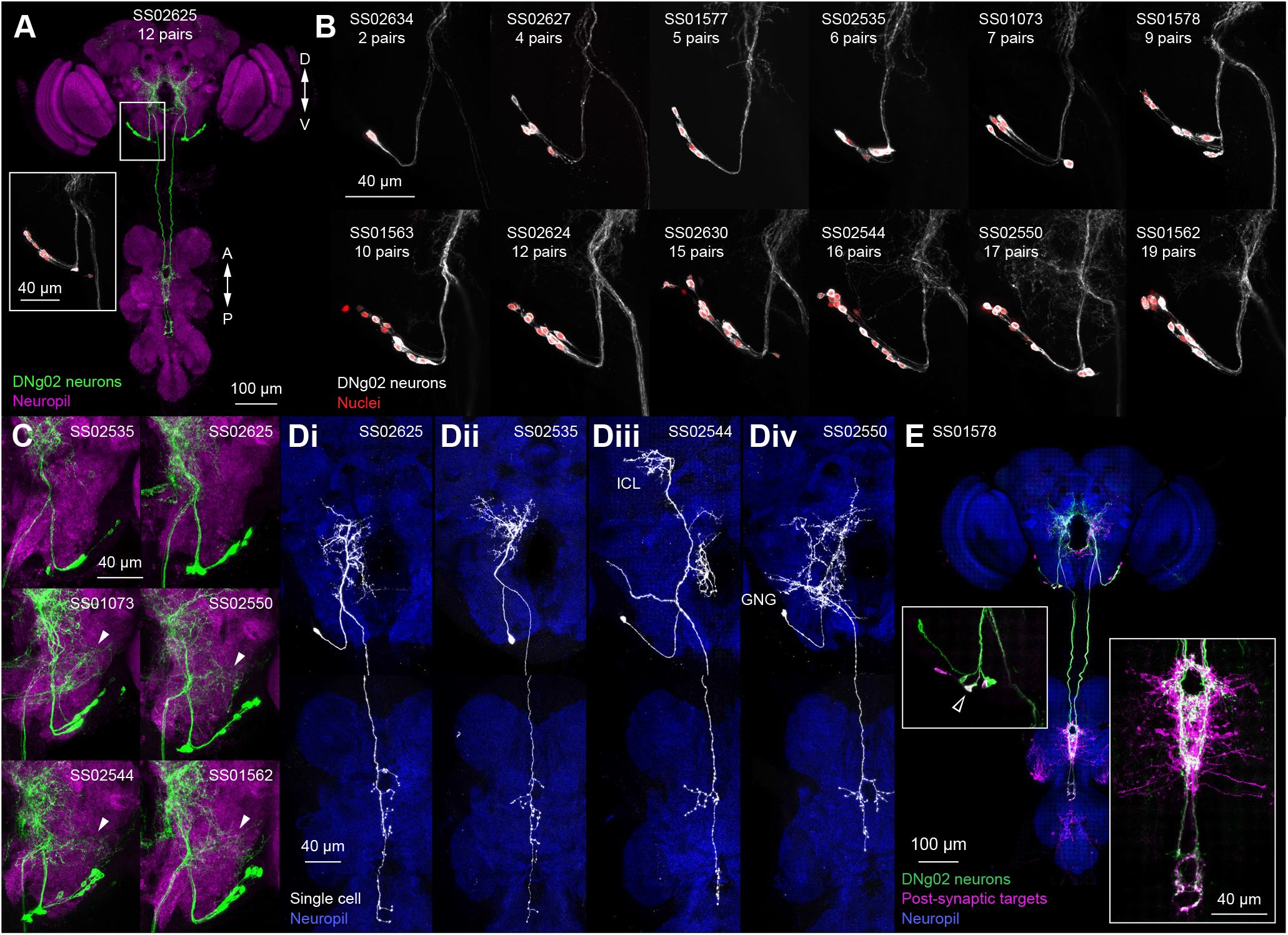
Morphology of DNg02 cells. (A-E) Confocal z-projections of DNg02 neurons in the fly central nervous system (CNS). Split-Gal4 ID numbers are indicated in each panel. (A) Expression pattern in an example driver line (SS02625) targeting DNg02 neurons in the full CNS. Green: membrane GFP; magenta: nc82. Inset: High magnification confocal z-projections of DNg02 cell bodies along the ventral ridge of the GNG. White: membrane GFP; red: nuclear localized DsRED. (B) Expression patterns represented as in (Figure 2A; inset) across the full set of split-Gal4 lines, sorted in ascending order based on the number of bilateral pairs in each line. (C) Expression patterns of six representative lines in the GNG. First row and bottom two rows display lines without and with GNG innervation (arrowheads), respectively. Green: membrane GFP; magenta: nc82. (D) Single cell labeling in the CNS using MCFO across representative split-Gal4 lines with variable innervation patterns: Type I DNg02 (Di), Type II DNg02 (Dii), DNg03 (Diii), and an unknown descending neuron (Div). White: MCFO – HA or FLAG; blue: nc82. ICL: inferior clamp; GNG: gnathal ganglion. (E) Postsynaptic targets of DNg02 neurons as revealed by trans-Tango in the full CNS. Green: membrane GFP in DNg02s; magenta: postsynaptic targets; blue: nc82. Left inset: DNg02 cell bodies. Coexpression of upstream and downstream labeling suggests reciprocal connectivity (empty arrowhead). Right inset: plexus of interneurons revealed in wing neuropil.

The morphology of DNg02 neurons have been characterized previously and were thought to be nearly homomorphic, possessing arborizations in inferior bridge (IB), inferior clamp (ICL), superior posterior slope (SPS), inferior posterior slope (IPS), as well as the GNG (Namiki et al., 2018; Namiki et al., 2022). Smooth processes in the IPS, SPS, and ICL suggest that these regions contain dendrites, whereas the presence of synaptotagmin-positive varicosities suggest that the IB and GNG contain output regions (Namiki et al., 2018). However, our new analysis identified subtle heterogeneities in the morphology of the cells targeted in the different driver lines. For example, some lines possess arbors that extend dorsally in the ICL, whereas others do not (Figure S2A). We observed another notable variation in the neurites within the GNG (arrowheads; Figure 2C), which was clearly observed in 4 of the 13 lines. To ascertain whether specific driver lines were enriched for particular DNg02 variants, we determined the morphology of individual cells by stochastically labeling neurons using the multicolor flip-out technique (Nern et al., 2015) (Figure 2D). For this analysis, we selected split-Gal4 drivers representing the variable arborization patterns observed across the lines. As innervation of the posterior slope in the brain, midline-crossing innervation of wing neuropil, and distal arbors in the haltere neuropil in the VNC appear to be the most characteristic DNg02 features, we used these as core criteria and identified two major variants within the set of cells identified as belonging to the DNg02 class (Figure 2Di-ii). The two variants innervate comparable neuropil regions in the brain with similar arborization patterns, but display a minor, but consistent, difference in the VNC. Whereas the DNg02 neurons described previously (Namiki et al., 2018; Namiki et al., 2022), which we refer to as Type I cells, exhibit a figure-of-eight shape in both the wing tectulum and haltere neuropil (Figure 2Di), a secondary variant, Type II cells, does not exhibit the clear figure-of-eight pattern (Figure 2Dii). Although the 13 drivers lines for DNg02 neurons were initially described as labeling members of a nearly homomorphic population of cells (Namiki et al., 2018; Namiki et al., 2022), our results indicate that different lines contain neurons that exhibit consistent differences in morphology, implying that there may be potential functional subdivisions within the population. It is unclear however, if the heterogeneity in cell morphology is substantial enough to warrant dividing DNg02 cells into two or more distinct DN types, or whether the variants are better viewed as subclasses of DNg02 cells.

We did, however, identify two neurons that while superficially resembling DNg02 cells, nevertheless warrant distinct classification (Figure 2D). Some lines contain a neuron that projects to the ICL, displays a lateral arbor in the GNG, does not innervate the posterior slope, and does not cross the midline in the VNC; rather, it projects laterally in the wing neuropil (Figure 2Diii). We believe this cell to be the previously identified DNg03 neuron (Namiki et al., 2018). Another cell type distinct from the DNg02s observed in some of the driver lines is a cell that prominently arborizes in the GNG and does not descend to the haltere neuropil (Figure 2Div). Given that lines SS01073, SS02550, SS02544, and SS01562 contain a combination of DNg02 neurons and other DNs innervating the GNG (Figure 2C), we focused subsequent functional analyses on the lines that specifically labeled the DNg02 variants (i.e., Type I and II DNg02 neurons).

To investigate connectivity downstream from the DNg02 population, we used the *trans*-Tango anterograde transsynaptic labeling system (Figure 2E) (Talay et al., 2017). Due to the density of staining, it is difficult to cleanly visualize most of the cell types downstream of the DNg02 neurons, but one notable exception is a population of interneurons with cell bodies situated in the ventral cell body rind of the VNC. These neurons project dorsally and splay out in a dense plexus of neurites in the wing neuropil adjacent to, and laterally of, the descending DNg02 fibers. We also found evidence for reciprocal connectivity within the DNg02 population. The *trans*-Tango preparations showed examples of likely DNg02 cells that exclusively contained a post-synaptic signal, as well as DNg02 neurons that contained both pre- and post-synaptic labels, consistent with monosynaptic connections among cells within the DNg02 population (inset; Figure 2E and Figure S2B). The presence of DNg02 cells with post-synaptic labeling indicates reciprocal connectivity within the population, although it is not possible to determine whether the connectivity includes connections across the midline from the *trans*-Tango results alone. We did not observe any motor neurons with post-synaptic labeling, although the possibility of false negatives using the *trans*-Tango method renders this observation difficult to interpret (Ni, 2021).

### Silencing DNg02 cells reduces the magnitude of the optomotor response

To investigate the role of DNg02 neurons in the optomotor response, we silenced the cells by selectively driving the inwardly rectifying potassium channel Kir2.1 (Baines et al., 2001) in the set of the nine sparse split-GAL4 driver lines that primarily targeted Type I and II DNg02 neurons. As controls, we also tested wild-type flies and flies in which we crossed UAS-Kir2.1 to a split-GAL4 line carrying empty vectors of the two GAL4 domains (SS03500). Silencing the DNg02 cells reduced the magnitude of the wing optomotor response, with a maximal reduction in ΔWBA from ~20° for the wild-type and empty vector control flies to ~10° for a driver line labeling 9 pairs of cells (Figure 3A). The reduction in ΔWBA during the optomotor response was roughly linear with the number of DNg02 cells silenced (slope = −0.6° cell^-1^, intercept = 20.6°) (Figure 3B). To test whether the observed trend could arise from chance, we performed a bootstrapping analysis in which we randomly sampled wing responses and numbers of cells silenced with 100,000 iterations. We found no iterations with a more extreme slope than the original dataset, giving a probability of zero that our experimental results arose from chance (Figure 3C). In addition, we performed a Wald test on the significance of the slope of the best fit line in Figure 3B, and found a p-value of 2 × 10^-8^ against the null hypothesis of zero slope. When the four driver lines labeling both DNg02 neurons and the neuron with arborization in the GNG were included in the analysis, we still found a statistically significant reduction in the magnitude of the optomotor response in proportion to the number of cell bodies silenced (Figure S3A-C). These results support the hypothesis that the DNg02 neurons regulate wingbeat amplitude via population coding. Both the outside wing (which increased in amplitude) and the inside wing (which decreased in amplitude) showed statistically significant reductions in response magnitude with the number of DNg02 neurons silenced, suggesting that DNg02 neurons are involved both in increases and decreases to wingbeat amplitude (Figure S4A-F). To determine how the silencing of cells altered the time course of the changes in ΔWBA, we again fit a Gaussian function to the derivative of the response curves (Figure 3D-E). We found a significant reduction in *ω_p_*, the maximum velocity of the change in the ΔWBA angle, a slight but significant increase in *t_p_*, the timing of the peak velocity, but no change in *d*, the duration of the response. This suggests that the decrease in the magnitude of the optomotor response is due to a reduction in the peak velocity of the change in ΔWBA, not in a decrease in the duration of the visual motion reflex.

**Figure 3.**
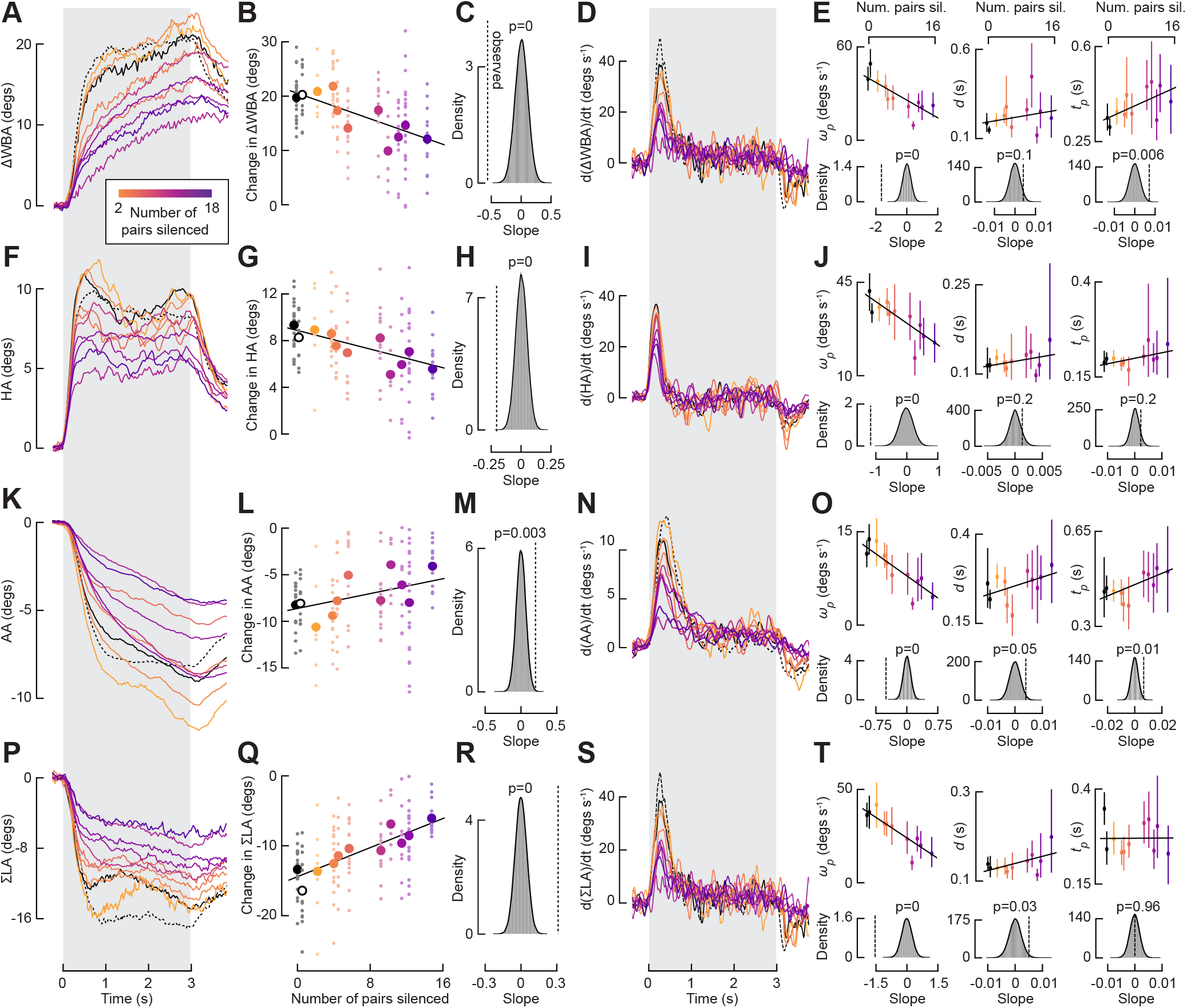
Silencing DNg02 neurons diminishes the magnitude of the optomotor response. (A) Averaged ΔWBA during the presentation of yaw motion for flies with DNg02 neurons silenced with Kir2.1 (sample sizes throughout figure: empty-split vector control: n = 19; wild-type control: n = 12; SS02634: n = 7; SS02627: n = 14; SS01577: n = 10; SS02535: n = 14; SS01578: n = 14; SS01563: n = 7; SS02625: n = 23; SS02624: n = 10; SS02630: n = 14). The number of pairs of cells silenced is indicated by the color of the trace, with the fewest pairs in yellow and the most in purple. The response of empty-vector control flies (SS03500) is shown with a solid, black line and the response of wild-type control flies is shown with a dotted, black line. The period of stimulus presentation is shown by the gray patch. (B) ΔWBA plotted against the number of DNg02 pairs silenced. As in Figure 3A, the number of pairs silenced is indicated by color. The two controls are offset slightly for visual clarity; the empty vector control is indicated by a solid circle and the wild-type control is indicated by an empty circle. Individual fly means are shown with small circles and grand means for a given driver line is shown with a large circle. A line is fit to individual fly means and plotted in black (r^2^ = 0.66, based on mean values). (C) Distribution of 100,000 bootstrapped slopes of lines of best fit with responses and numbers of cells silenced randomly sampled with replacement. The dotted, black line indicates the observed slope of the original dataset. The p-value is the proportion of resampled slopes that result in a more extreme slope than observed. (D) Time derivative of the ΔWBA data shown in Figure 3A, calculated using a Savitzky-Golay filter. (E) Optomotor response is reduced due to a decrease in angular velocity, not a decrease in the duration of movement. Top: As in Figure 3B, for the parameters from fitting a Gaussian function to the data in Figure 3D (from left to right, r^2^ = 0.67, 0.08, 0.34, based on mean values). Bottom: As in Figure 3C, for the Gaussian function parameters. (F) As in Figure 3A, for head optomotor response. (G) As in Figure 3B, for head optomotor response (r^2^ = 0.66, based on mean values). (H) As in Figure 3C, for head optomotor response. (I) As in Figure 3D, for head optomotor response. (J) As in Figure 3E, for head optomotor response (top row, from left to right, r^2^ = 0.65, 0.16, 0.33, based on mean values). (K) As in Figure 3A, for abdomen optomotor response. (L) As in Figure 3B, for abdomen optomotor response (r^2^ = 0.41, based on mean values). (M) As in Figure 3C, for abdomen optomotor response. (N) As in Figure 3D, for abdomen optomotor response. (O) As in Figure 3E, for abdomen optomotor response (top row, from left to right, r^2^ = 0.78, 0.21, 0.46, based on mean values). (P) As in Figure 3A, for leg optomotor response. (Q) As in Figure 3B, for leg optomotor response (r^2^ = 0.85, based on mean values). (R) As in Figure 3C, for leg optomotor response. (S) As in Figure 3D, for leg optomotor response. (T) As in Figure 3E, for leg optomotor response (top row, from left to right, r^2^ = 0.79, 0.31, 0.004, based on mean values).

As with the wing response, we saw a significant reduction in the head, abdomen, and leg responses upon silencing DNg02 neurons (Figure 3F-T; Figure S4G-L). Again, the head, abdomen, and leg optomotor responses appear to be diminished due to a reduction in the angular velocity of the motor response, not a decrease in its duration. Whereas DNg02 neurons directly innervate regions of the VNC associated with flight and other wing-related behaviors, the cells do not make direct projections to the neck, leg, or abdomen neuropils. While this projection pattern does not preclude the existence of connections to motor circuits via a poly-synaptic pathway, it is also possible that the reduction in head, leg, and abdomen optomotor response is mediated indirectly via a decrease in the strength of mechanosensory reflexes driven by the changes in wing motion caused by silencing DNg02 cells. When the four driver lines labeling both DNg02 neurons and the neuron with arborization in the GNG were included in the analysis, we still observed significant reductions in the magnitudes of the head and leg responses, although the effect on the abdomen response was not significant (Figure S3D-L). The number of DNg02 pairs silenced did not alter the baseline wingbeat amplitude during flight (Figure S5); therefore, such an effect could not explain the reduction in any of the visually induced motor responses we observed.

### DNg02 activity correlates to motor output and is consistent across morphological variants

Our silencing experiments indicate that DNg02 neurons are required for the optomotor response to reach its full magnitude. To investigate how the DNg02 cells respond to the visual optomotor stimulus, we performed two-photon functional imaging in tethered flying flies using GCaMP8m (Zhang et al., 2021) as an activity indicator (Figure 4A-B). We selected driver lines that labeled either only the Type II DNg02 variant (that is, lines without a clear figure-of-eight shape within the haltere neuropil), or lines that likely labeled both Type I and Type II DNg02 neurons (i.e., lines that contained at least some cells with the clear figure-of-eight feature), in order to attempt to identify functional roles of the two variants. During these experiments, flies often stop flying, in which case we would try to re-initiate flight with a puff of air. We made use of these occasional flight stops (and subsequent puff-induced starts) to record the activity of DNg02 neurons during the transitions between flight and quiescence (Figure 4C). We found a higher level of activity during flight than quiescence, a trend consistent with other cells in the brain (Maimon, et al., 2010).

**Figure 4.**
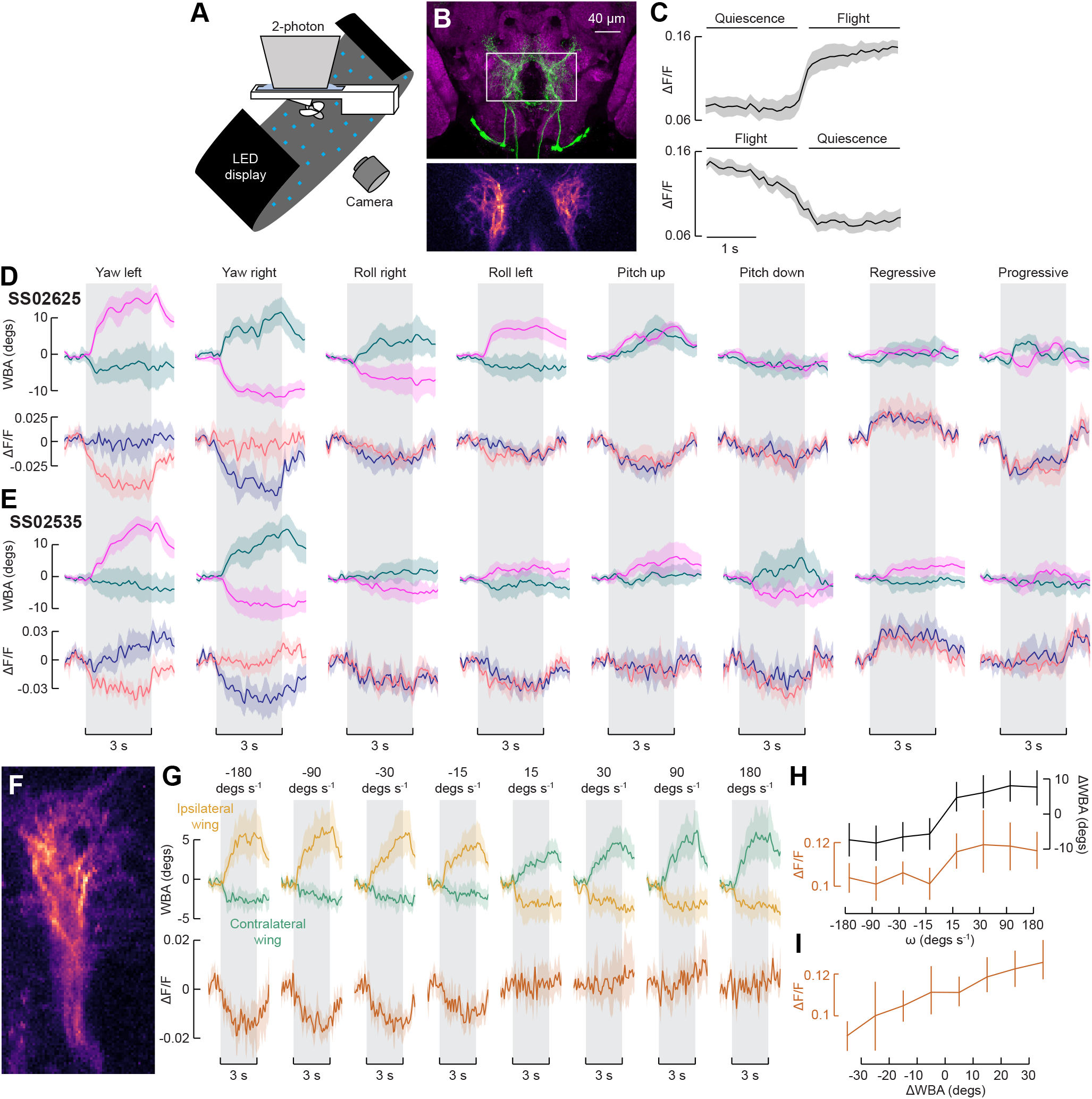
DNg02 activity correlates with visually-elicited motor responses. (A) Schematic showing the experimental set-up used for two-photon functional imaging. A fly is tethered to a flight stage under the microscope at the center of an array of LEDs. A camera tracks wing position in real-time using a machine vision system. (B) We imaged a region, outlined in white, capturing DNg02 branches bilaterally. Bottom: standard deviation of GCaMP8m fluorescence over time. The 10% most variable pixels are used to define a region of interest within which we measure changes in fluorescence. See Methods for details. (C) Average fluorescence as the fly transitions between quiescence and flight, agnostic to visual stimuli (n = 7 flies, 198 transitions from quiescence to flight, 188 transitions from flight to quiescence). Bilateral fluorescence increases at the onset of flight and decreases at the offset. (D) Averaged wing and fluorescence responses to different patterns of optic flow for SS02625, a driver line labeling both Type I and Type II DNg02 variants (n = 9 flies). Top row shows baseline-subtracted left (teal) and right (pink) wingbeat amplitudes; bottom row shows baseline-subtracted left (blue) and right (salmon) DNg02 activity. Data are presented as mean (solid line) and a bootstrapped 95% CI for the mean (shaded area). The 3 s stimulus period is shown by the gray patch. (E) As in Figure 4D, for SS02535, a driver line labeling exclusively Type II DNg02 neurons (n = 11 flies). (F) Standard deviation of GCaMP8m fluorescence for an example of unilateral imaging. Cells on the right side of the brain are shown. The driver line used for these experiments, SS02625, labels DNg02 neurons of both Type I and Type II morphology. (G) Averaged wing and fluorescence responses to yaw motion at a range of angular velocities (n = 14 flies). Ipsilateral and contralateral wingbeat amplitudes shown on the top row; DNg02 fluorescence shown on the bottom. Plotting conventions as in Figure 4D. (H) Mean fluorescence and ΔWBA (ipsilateral amplitude subtracted from contralateral) across stimulus angular velocities (n = 14 flies). Mean is taken over the final second of stimulus presentation (i.e., from t = 2 to 3 s). Error bars show bootstrapped 95% CI. (I) Mean fluorescence across as a function of ΔWBA, with data parsed within 8, 10°ΔWBA bins. The plot incorporates data collected across the full range of visual stimulus conditions. Error bars show bootstrapped 95% CI.

To probe the role of the DNg02 neurons in flight stabilization and any functional variations between the Type I and II variants, we presented wide-field visual stimuli simulating the optic flow flies would experience during pitch, roll, and yaw rotations of the body, as well as progressive and regressive translations (Figure 4D-E). These stimuli were selected to induce a diverse array of flight behaviors. Whereas roll and yaw stimuli induce asymmetric wingbeat responses, pitch, regressive, and progressive stimuli induce symmetric responses (Götz, 1983). Presentations of optomotor stimuli were interspersed with epochs of a static starfield to return wing kinematics to baseline. Broadly, our results recapitulate previously reported responses in DNg02 neurons to these visual stimuli in one specific driver line (Namiki et al., 2022), with some notable exceptions (Figure 4D-E). Previous work reported robust increases and decreases in DNg02 activity correlated with contralateral wing motion during the presentation of yaw motion (e.g., in response to leftward drifting stimulus, the right wing increased in amplitude as the left DNg02 increased in activity, while the left wing decreased in amplitude as the right DNg02 decreased in activity). In contrast, we observed only weak increases in DNg02 activity during the presentation of yaw motion, although we did observe robust decreases in activity. We believe that this may be due, in part, to the different calcium indicator used; the prior work used GCaMP6f while we used GCaMP8m. Furthermore, whereas previous work reported asymmetric neuronal responses without behavioral responses to roll motion, we observed asymmetric behavioral responses, as quantified by changes in wing kinematics, and weak symmetric decreases in neuronal activity. The differences in our results could also be due to imaging a slightly different region of the brain, which could result in recording the activity of different DNg02 cells than in the previous study. Regardless, the broad results are the same as previously reported; that is, DNg02 activity correlates with contralateral wingbeat amplitude and DNg02 cells can respond independently on the left and right sides of the brain. We further note that the highest magnitude neuronal responses are induced by yaw, regressive, and progressive motion, all of which consist primarily of patterns of horizontal optic flow. We repeated the experiments with driver lines labeling both Type I and Type II DNg02 neurons (Figure 4D, S6A), exclusively Type II cells (Figure 4E, S6B), and a line labeling Type II neurons and neurons with GNG innervation (Figure S6C) and found that the responses are the same regardless of the driver line imaged.

Because the results of both the silencing screen we present here and the activation screen in a prior study (Namiki et al., 2022) suggest that the DNg02 cells likely operate via population coding, we performed a further set of functional imaging experiments to determine if we could find evidence for stimulus-dependent recruitment, such that the number of cell activated increases with the strength of the visual stimulus. Given that the strongest responses were to yaw motion, we presented these patterns of optic flow across a range of stimulus angular velocities, in order to induce different magnitude neuronal and behavioral responses. To maximize our ability to resolve individual cells, we only imaged the DNg02 cells on one side of the brain (Figure 4F). However, we were not able to resolve individual cells to specifically test whether different neurons became active at different strengths of the visual stimulus or were correlated with different magnitudes of the motor response. The driver line we tested (SS02625) labels 12 pairs of neurons, but it is possible that a line targeting fewer pairs would allow us to distinguish individual cells. As has been reported previously (Namiki et al., 2022), DNg02 activity appears to be correlated with contralateral wingbeat amplitude and anticorrelated with ipsilateral wingbeat amplitude relative to the side of the brain on which we imaged (Figure 4G). Plotting the ΔWBA (here, ipsilateral wingbeat amplitude subtracted from contralateral) and the fluorescence signal within the ROI (ΔF/F) against the angular velocity of the pattern presented (Figure 4H) indicates that both responses saturate at velocities of >15° s^-1^. However, when we directly plotted the fluorescence signal against ΔWBA (independent of the angular velocity at which the data were collected), we found a strongly linear relationship (Figure 4I), suggesting that DNg02 activity is tightly coupled to the motor response. While the silencing experiments and previously reported activation experiments (Namiki et al., 2022) suggest that there is a causal relationship between DNg02 activity and wing motion, this coupling could also arise in part from ascending feedback signals from neurons in the VNC.

### Unilateral DNg02 activation induces bilaterally symmetric behavioral responses

Both the silencing screen and the two-photon functional imaging results are consistent with the hypothesis that DNg02 neurons regulate contralateral wingbeat amplitude. If so, unilateral DNg02 activation should induce strong steering maneuvers, in which the contralateral wing increases its amplitude upon activation, and the ipsilateral wingbeat amplitude either decreases (in the case in which DNg02 neurons inhibit motor output on the ipsilateral side) or does not change (in the case in which DNg02 neurons have effect on ipsilateral wing motion). To test this hypothesis, we targeted optogenetic stimulation of DNg02 neurons on one side of the central nervous system (CNS) via a genetic strategy used previously for lobula columnar neurons (Wu et al., 2016). This approach leverages the modular organization of insect nervous systems; neurons of the CNS descend from individual neural progenitors called neuroblasts, each of which produces a unique lineage of developmentally related cells (Spindler and Hartenstein, 2010). Using this technique, we could create animals expressing Chrimson-Venus in DNg02 neurons unilaterally (Figure 5A); we also generated animals with bilateral or no labeling (Figure 5B-C), providing useful controls with identical genetic backgrounds and experimental rearing conditions. We scored the expression pattern in the brain of each fly after the conclusion of an experiment.

**Figure 5.**
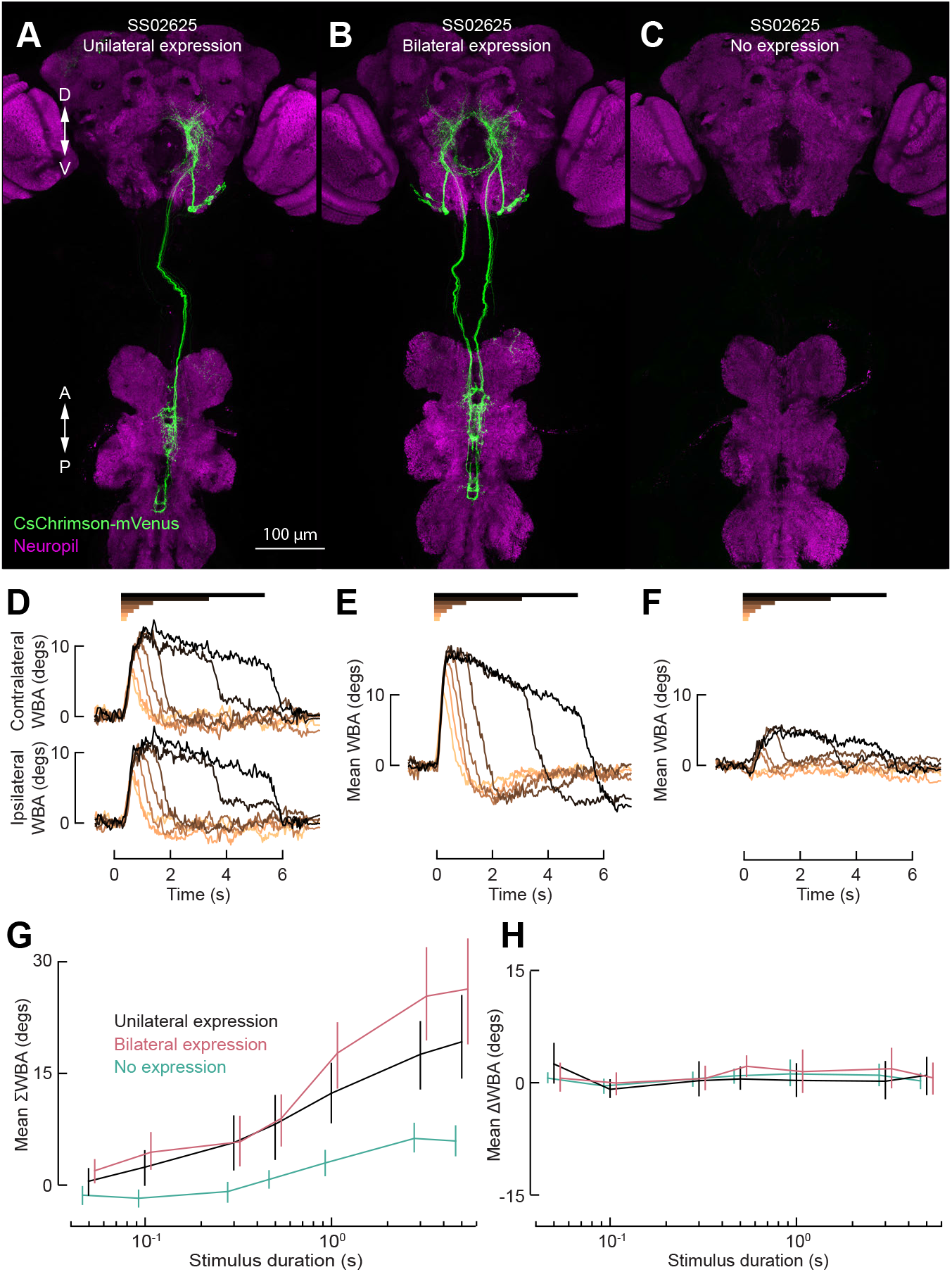
Unilateral DNg02 activation induces bilaterally symmetric changes to wing kinematics. (A-C) Confocal z-projections of the fly central nervous system. Flp-recombinase under heat shock control coupled with 20XUAS-FRT>-dSTOP-FRT>-CsChrimson-mVenus enables temperature dependent, stochastic expression of CsChrimson-Venus (green) in DNg02 neurons unilaterally (A), bilaterally (B), or neither (C). Magenta: nc82. (D) Baseline-subtracted contralateral and ipsilateral WBA responses of flies with unilateral expression of CsChrimson to a red light stimulus of variable duration (0.05, 0.1, 0.3, 0.5, 1, 3, or 5 s, indicated by the horizontal bars above) (n = 8 flies). Flies are presented with a grating visual pattern in closed loop with their left minus right WBA. (E) Baseline-subtracted mean WBA responses of flies with bilateral expression of CsChrimson (n = 14 flies). Plotting conventions as in Figure 5D. (F) As in Figure 5E, for flies with no expression of CsChrimson (n = 23 flies). (G) Mean ΣWBA against duration of stimulus. Expression pattern is indicated by color, with unilateral expression in black, bilateral in pink, and no expression in green. Vertical bars indicate a bootstrapped 95% CI. Bilateral and no expression traces are offset slightly for visual clarity. (H) As in Figure 5J, for ΔWBA.

Upon activation, flies with cells labeled bilaterally exhibit strong increases in the bilateral mean of wingbeat amplitude, consistent with previous experiments (Namiki et al., 2022) (Figure 5E, G). Control flies in which no cells expressed CsChrimson exhibited no changes in mean WBA, at least for stimulus duration less than 300 ms (Fig 5F, G). At longer stimulus durations, control flies did show small increases in mean WBA, which we interpret to be an artifact induced either by a behavioral response to the 617 nm light or the heat it generated. In the case of unilateral DNg02 activation, we expected to observe a strong decrease in ΔWBA (here, ipsilateral – contralateral WBA) in response due to an increase in wingbeat amplitude on the side contralateral to the stimulated cells. In contrast, however, we observed comparable increases in both contralateral and ipsilateral wingbeat amplitude and thus a large mean WBA response (Figure 5D, G). Further, in all three cases (unilateral, bilateral, or no activation), the ΔWBA response was very close to zero in response to activation regardless of stimulus duration (Figure 5H); thus, wing kinematics remained bilaterally symmetric regardless of whether the activation was symmetric or asymmetric. The only difference between unilateral and bilateral activation was that the resulting mean WBA response was higher with bilateral activation (Figure 5G), which is consistent with more cells being activated under that condition (i.e., the DNg02 neurons on both the left and right sides of the brain are activated). We repeated the experiments at lower stimulus durations and across different driver lines and found the same result: regardless of stimulus duration or number of cells labeled, the resulting behavior was a bilaterally symmetric increase in wingbeat amplitude (Figure S7). These results suggest that unilateral optogenetic recruitment of DNg02 activity drives bilaterally symmetric motor output, although the precise mechanism (e.g., reciprocal connectivity within the population, symmetric bilateral downstream connectivity to motor neurons) is unknown.

## Discussion

In this paper, we examined the role of a population of descending neurons in the optomotor response of flying *Drosophila*. The optomotor response consists of coordinated movements of different appendages, including changes in wing kinematics, head position, and deflections of the abdomen and hind legs. Prior work based on anatomy and optogenetic activation identified the DNg02 neurons as likely candidates for regulating mechanical power and mediating steady-state flight control by transmitting optic flow from the brain to motor centers in the VNC. In this paper, we further tested this hypothesis through a combination of anatomy, genetic silencing, unilateral optogenetic activation, and functional imaging. When DNg02 neurons were inactivated via chronic expression of the inward-rectifying potassium channel Kir2.1, the magnitude of the optomotor response was diminished in proportion to the numbers of cells silenced, up to a value of ~50% compared to the responses of wild-type or empty-vector control flies (Figure 3A,B). It is possible that DNg02 neurons provide direct input to flight motor neurons or local pre-motor interneurons, as the cells project to the wing neuropil. However, we also observed a similar reduction in the optomotor responses of the head (Figure 3F, G), abdomen (Figure 3K,L), and legs (Figure 3P,Q), although these reflexes are less likely to be driven directly by DNg02 cells as they do not project to the relevant motor neuropils. It is unclear whether the reductions in these responses are due to polysynaptic connections from the DNg02 cells to neck, leg, and abdomen motor neurons, or alternatively, through changes in reflexive pathways that are reduced by the diminution of the wing response. For example, if mechanosensory feedback from wing mechanoreceptors is partially responsible for eliciting movements of the neck, legs, and abdomen, these reflexes might be attenuated by the reduction in the magnitude of the wing response. Further experiments will be necessary to test between these two alternatives.

There are several non-mutually exclusive explanations for why genetic silencing did not result in complete abolishment of the optomotor response. First, even with the driver line labeling the highest number of cells, we may not have been able to silence the complete set of DNg02 neurons. Second, it is possible that Kir2.1 expression does not completely abolish the ability of these cells to conduct spikes to the VNC. Finally, there may be additional, as of yet unknown, classes of DNs that are involved in the optomotor response besides DNg02. Give that unilateral activation of the DNg02 cells did not elicit asymmetrical changes in wing motion (Figure 5), this last hypothesis is particularly relevant for developing a comprehensive explanation for all the results in our study.

We conducted additional experiments that sought to illuminate the possible role of different morphological types of DNg02 neurons in the optomotor response. Two-photon imaging experiments suggest that morphological differences within the DNg02 population do not appear to correspond to functional subpopulations, at least with respect to the responses elicited by the set of wide-field patterns of optic flow that we presented. It is possible, however, that our stimulus set did not include patterns of visual motion (or other sensory stimuli) that would illuminate functional differences among the different DNg02 types. The cells may also diversify functionally within the VNC, such that all DNg02 cells exhibit the same patterns of activity but have differing effects on the motor system. If that is true, however, we were not able to detect such differences by monitoring wingbeat amplitude, but different DNg02 types might elicit changes in other features of wing motion that are not detected by our machine vision system (e.g., changes in stroke deviation or angle of attack). The DNg02 cells appeared to have the strongest responses to patterns of visual motion in the horizontal plane (i.e., yaw, regressive, and progressive motion) and neurons on the left and right side of the nervous system can be active independently from each other. Responses to yaw motion over a range of rotational velocities suggest that DNg02 activity is tightly coupled to the motor response of the wings. While this coupling could reflect efferent copy or ascending reafferent sensory input to the DNg02 cells, the activation and silencing experiments indicate a causal relationship between DNg02 activity and motor output, although the role of feedforward and feedback mechanisms are not mutually exclusive.

Given the result of our silencing experiments, it is noteworthy that unilateral optogenetic activation of DNg02 neurons did not elicit bilaterally asymmetric changes in wing motion similar to those elicited by a visual yaws stimulus, but rather evoked symmetric increases in wingbeat amplitude comparable to those elicited by bilateral activation. Whereas the silencing and functional imaging experiments suggest that DNg02 neurons regulate contralateral wingbeat amplitudes via population coding, the unilateral activation results do not support this hypothesis. There are several possible explanations for this discrepancy, which are not mutually exclusive. First, if the reciprocal connectivity within the DNg02 population identified by our *trans*-Tango experiments includes connections among DNg02 cells on the left and right side of the body, unilateral activation might recruit other members of the population across the midline. Unfortunately, our *trans*-Tango results do not distinguish between ipsilateral and contralateral connections, so we are not able to further evaluate this possibility. The fact that the DNg02 cells do display bilaterally asymmetric responses when presented with visual yaw motion (Figure 4), would suggest that the coupling among the cells is not so strong so as to lock bilateral pairs of neurons within the population to the same level of activity.

Second, it is possible that to induce asymmetric motor output, DNg02 activity must be in opposition across the midline such that cells on one side increase in activity while those on the other side decrease in activity. The large change in the level of activity at the onset of flight (Figure 4C) is relevant to this hypothesis, because it indicates that the neurons might encode visual motion via both increases and decreases in changes in the background level of activity. Indeed, the interpretation of unilateral DNg02 activation is unclear given our functional imaging experiments, as we never recorded increases in cell activity on one side with no changes on the other side of the brain during our functional imaging experiments (Figure 4D). We did observe decreases in activity on one side and no changes on the other, but we are unable to elicit this pattern of activity via optogenetic methods. Although optogenetic silencing channels are available (Mohammad et al., 2017), preliminary experiments indicated that the strong behavioral artifacts elicited in flying flies by the presentation of the blue light required for GtACR activation render the results uninterpretable.

Third, it is possible that DNg02 neurons project bilaterally to flight motor neurons on both the contralateral and ipsilateral side. As described in the Introduction, two distinct types of flight muscles regulate wing motion in flies: large asynchronous power muscles, which provide power to the wings indirectly via their action on the scutellar lever arm and the anterior notal wing process (Boettiger and Furshpan, 1952; Deora et al., 2015), and the tiny synchronous steering muscles, which generate subtle changes in wingbeat kinematics via their directly insertions on elements within the wing hinge (Tu and Dickinson, 1997; Lehmann et al., 2013; Lindsay et al., 2017). It is quite possible that both muscle systems are involved in the optomotor response, because sustained changes in wing motion require regulation both of power output and wing kinematics. In particular, the mechanical power required to flap a wing back and forth scales roughly with (stroke amplitude)^3^ (Ellington, 1984). Thus, an increase in wing stroke amplitude would require an increase in mechanical power, where as a decrease in stroke amplitude could still be sustained with decrease in mechanical power. The steering muscles are not thought to contribute to the positive power required to flap the wings; rather they perform negative work, but function as controllable dynamic springs to regulate the mechanics of the wing hinge (Tu and Dickinson, 1994). Namiki and coworkers (2022) showed that DNg02 activation increases the total mechanical power output of the flight system, suggesting that these neurons provide a strong drive to the motor neurons of the asynchronous power muscles. Our results on unilateral activation support this conclusion, but the question remains why silencing these cells results in a reduction of the optomotor response that we observed.

Interpreting the roles of the power and steering and power muscles during a sustained optomotor response is complicated by several factors. First, because of the asymmetry in the kinematic response, the power requirements for one wing will increase while that for the other wing decreases. In our experiments, we recorded an increase in stroke amplitude of the outside wing of ~13° and a decrease of the inside wing of ~7°, which would require a net increase in power delivered to the entire system. However, if the mechanical design of the thorax and wing hinge is such that the motion of the scutellar lever arms and anterior notal wing processes are linked across the two sides of the body, is it even possible for the DLMs and DVMs to deliver different amounts of mechanical power to the two wings? Second, if flies were optimized to minimize energy consumption, the power delivered to each wing would be exactly that required to sustain the inertial and aerodynamic power required to flap it back and forth, but this need not be the case. The insect flight system in known to be quite inefficient, with losses in mechanical power estimated to be ~90% (Lehmann and Dickinson, 1997). Even if this value is not precise, the fact remains that flies could generate the changes in wing kinematics required for the optomotor response simply by slightly increasing the efficiency of one wing hinge while reducing the efficiency of the other—with no net change in total mechanical power required. These putative changes in efficiency might be achieved via the action of the steering muscles.

Because measuring the mechanical efficiency of an individual wing hinge is a task that far exceeds current experimental means, this speculative topic is best set aside for future investigation. Instead, it is worth considering the possibility that the fly is indeed capable of delivering different amounts of mechanical power to the two wings, despite the fact that the architecture of the thorax, particularly the structure of the scutellum and sculler lever arms, would seem to preclude this. It is noteworthy, however, that Lehmann and co-workers (2013) measured bilateral differences in the Ca^2+^ signals within both the DLMs and DVMs during presentation of optomotor stimuli, strongly suggesting that the left and right sets of muscles can be activated asymmetrically. Could such patterns translate into a differential mechanical drive of the left and right wings? While it is true that the scutellar lever arms appear to be rigidly linked across the two side of the thorax (Deora et al., 2015), this does not entirely exclude the possibility that this mechanical system could exhibit a bilateral ‘wobble’, such that the oscillatory trajectory of the posterior notal wing process (the tip of the scutellar lever arm that contacts the second axillary sclerite) might be larger on one side of the fly than the other, thus allowing differential drive to the two wings. According to this hypothesis, this bilateral asymmetry might be regulated via careful coordination of steering and power muscle activity, the latter of which involves the action of the DNg02 cells. Furthermore, whereas the mechanical role of the scutellar lever arms in coordinating the motion of the two wings has been well studied (Boettiger and Furshpan, 1952; Deora et al., 2015), little is known about the anterior notal wing process, which is thought to serve as the other primary means of transmitting the strains of the power muscles to the wing. Due to its more flexible attachment to the notum (Boettiger and Furshpan, 1952; Miyan and Ewing, 1984; Pringle, 1957) and the more lateral position of the DVMs, it is possible that this structure more easily permits differential actuation of the two wing hinges than the scutellar lever arms, which are actuated by the more medially positioned DLMs.

In summary, we propose that flies might be able to differentially regulate the mechanical power delivered to the two wings by the DLMs and DVMs. This active process might be mediated by a combination of asymmetrical activation of the power muscles themselves (as observed by Lehmman et al., 2013) and asymmetrical regulation of the left and right hinge mechanics by the steering muscle system, generating a bilateral wobble in the motion of scutellar lever arm system discussed above. This mechanism could explain why silencing the DNg02 cells would reduce the magnitude of the optomotor response (because it reduces the total mechanical power going to the left and right wings), but that unilateral activation of DNg02 neurons is alone insufficient to recapitulate the large bilateral changes in wing motion characteristic of the optomotor response (because regulation of hinge mechanics requires the action of the steering muscles). Under this hypothesis, alternative descending pathways would be responsible for transmitting commands to the steering motor neurons, which are bilaterally uncoupled and can act independently. Hopefully, this somewhat complex hypothesis will eventually be testable, once data from a complete *Drosophila* nervous system connectome (i.e., brain and VNC) are available.

In this paper, we have demonstrated that the DNg02 neurons are required for the optomotor responses of the wings, head, abdomen, and legs to reach their full magnitude. Given that unilateral activation of the DNg02 population results in bilaterally symmetric motor responses, the precise mechanism of control of the population over wing kinematics remains elusive, but we suspect a full explanation requires a more nuanced appreciation for the role of power muscles in flight maneuvers, and a deeper understanding of how steering muscles regulate the transmission of mechanical power to the two wings. Regardless, identification of the DNg02 neurons as an important component in the optomotor response provides a convenient entry point into a wide array of inquiries regarding the descending control of flight behavior in insects.

## ACKNOWLEDGMENTS

We would like to thank all members of our lab for helpful discussions. In particular, we thank Will Dickson for building the laser cutter used for the functional imaging experiments. We also thank the Parker laboratory for confocal microscope access. Research reported in this publication was supported by the National Institute of Neurological Disorders and Stroke of the National Institutes of Health under Award U19NS104655.

## AUTHOR CONTRIBUTIONS

(Following CRediT taxonomy): conceptualization—E.H.P., J.J.O., and M.H.D.; investigation— E.H.P. and J.J.O.; resources—J.J.O.; visualization—E.H.P. and J.J.O.; methodology—E.H.P., J.J.O.; software—E.H.P., formal analysis—E.H.P.; validation—E.H.P., writing original draft— E.H.P., M.H.D.; writing review and editing—E.H.P., J.J.O., and M.H.D.; J.J.O.; data curation— E.H.P.; funding acquisition—M.H.D.; supervision—M.H.D.; project administration—M.H.D.

## METHODS

### KEY RESOURCES TABLE

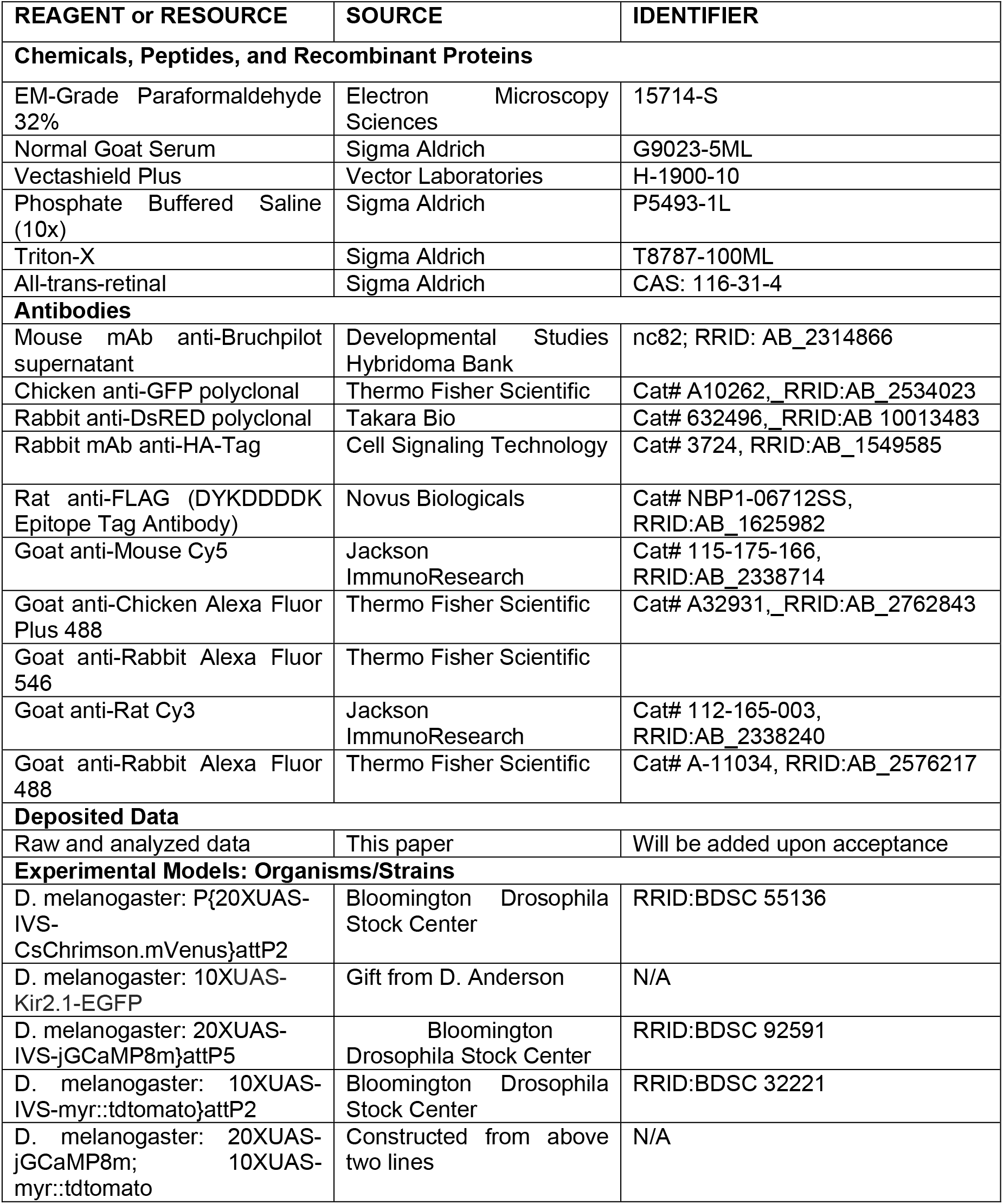
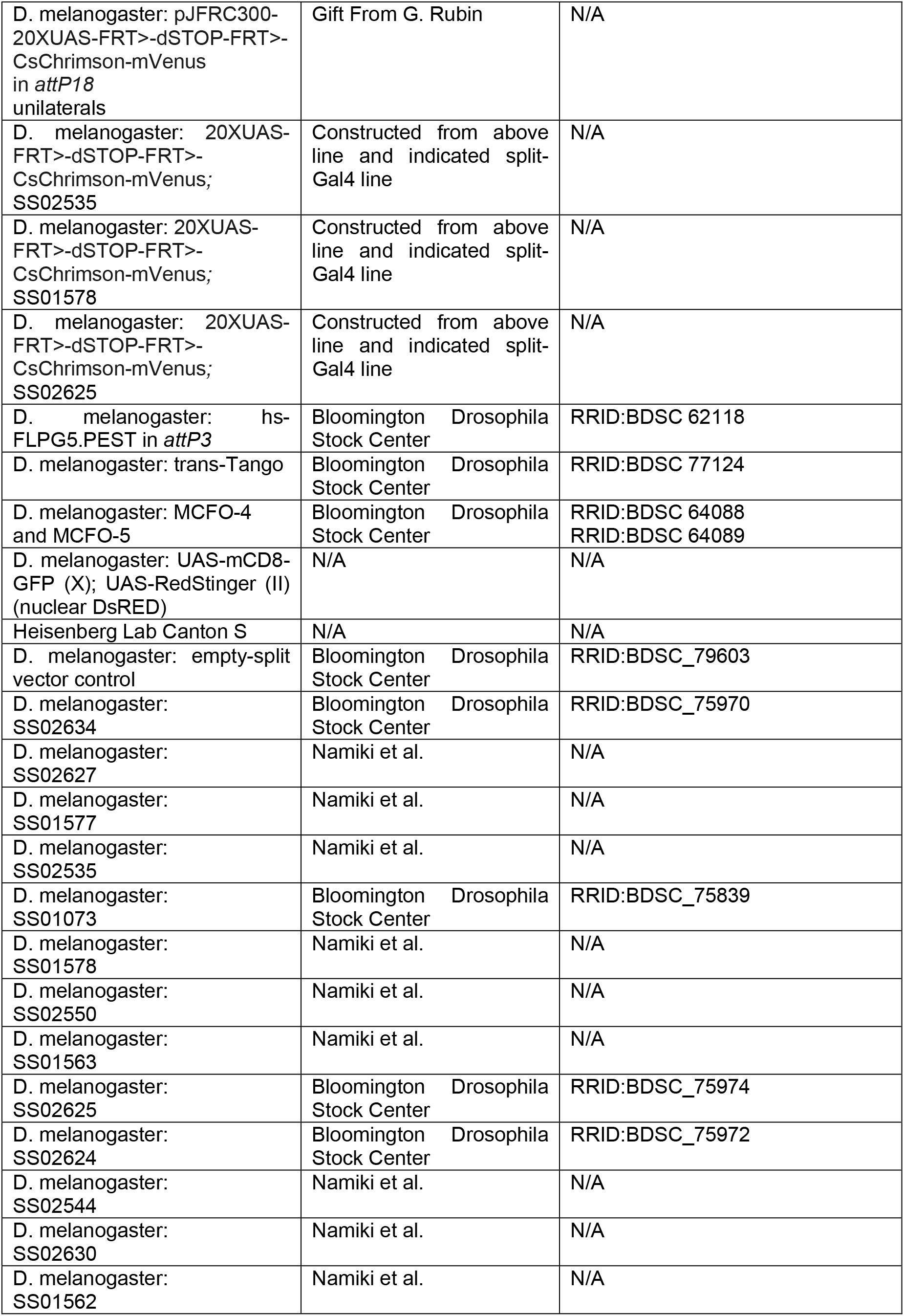
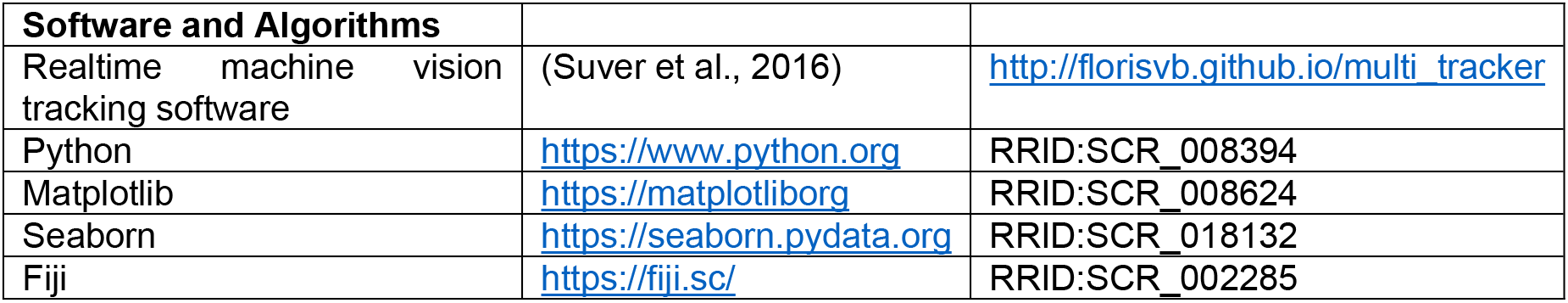

### CONTACT FOR REAGENT AND RESOURCE SHARING

Further information and requests for resources and reagents should be directed to and will be fulfilled by the Lead Contact, Michael H. Dickinson.

### EXPERIMENTAL MODEL AND SUBJECT DETAILS

All experiments were conducted on 2-to-5 day old female *Drosophila melanogaster* reared at 25°C on a 12:12 hr light-dark cycle unless otherwise specified. For the optogenetic activation experiments, we reared the flies on standard cornmeal fly food containing 0.2 mM trans-Retinal (ATR) (Sigma-Aldrich) and transferred flies 0-2 days after eclosion onto standard cornmeal fly food with 0.4 mM ATR in dark conditions. For all other experiments, flies were reared on standard cornmeal fly food. All standard cornmeal food was supplemented with additional yeast. For the silencing experiments, we used flies resulting from a cross of each split-GAL4 line with 10XUAS-Kir2.1-EGFP. For functional imaging experiments, we used flies resulting from a cross of the split-GAL4 lines with 20XUAS-jGCaMP8m; 10XUAS-myr::tdtomato. Similarly for expression analysis, we crossed split-Gal4 lines to UAS-mCD8-GFP; UAS-RedStinger. To trace post synaptic partners of DNg02 neurons, specified split-Gal4 lines were cross to trans-Tango; F1 progeny were stored in 18°C for ~3 weeks before dissection and processing.

For unilateral activation experiments, we used a “Flp-out” construct with UAS sequences upstream and Chrimson-Venus as the downstream effector. Flp recombinase expression under heat shock control during development can stochastically elicit temporal excision of transcriptional stop sequences in the parent neuroblast of DNg02 neurons, resulting in the heritable, permissive conformation of Chrimson-Venus under DNg02 split-Gal4 control. For these experiments, we used flies resulting from a cross of pJFRC300-20XUAS-FRT>-dSTOP-FRT>-CsChrimson-mVenus in *attP18* on (X) combined with specified split-Gal4 hemidrivers on (II) and (III) chromosomes with hs-FLPG5.PEST in *attP3* (X). Temperature-shift dependent CsChrimson-Venus-labeled DNg02 clones were induced by heat shocking 0-12 hour old embryos in a bottle for 60-90 min in a 37°C water bath. F1 progeny were returned to standard rearing conditions (25°C) until eclosion; adults were transferred to 0.4mM ATR food for ~3 days in dark conditions until experimentation. For single cell analysis using the multicolor flip out method (Nern et al., 2015), we used flies resulting from the cross of a specified Gal4 driver to MCFO-4 or MCFO-5 depending on the cell density of the driver (MCFO-4 and MCFO-5 for <10 and >10 pairs of neurons, respectively). For bilateral activation experiments, we used flies reared in optogenetic conditions resulting from a cross of male flies from the split-GAL4 line SS02625 with *UAS-CsChrimson* female virgin flies.

### QUANTIFICATION AND STATISTICAL ANALYSIS

All experiments were analyzed with custom scripts written in Python. Variance across individuals is quantified as the bootstrapped 95% confidence interval using Seaborn statistical functions, with 1,000 bootstrap iterations. We performed a bootstrapping analysis to assess whether the results in Figures 3, S3, S4, and S5 were significant. In each iteration, we randomly sampled with replacement from both the behavioral metric dataset (e.g., ΔWBA) and the number of pairs silenced, such that responses were randomly paired to numbers of pairs. Then, we fit a line of best fit to the bootstrapped dataset. The p-value was then calculated as:

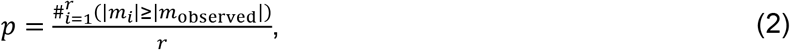

where *r* = 100,000 is the number of iterations, *m_i_* is the slope of the *i*^th^ iteration, and *m_observed_* is the observed slope. The p-value is therefore the fraction of iterations yielding a more extreme slope than observed (MacKay). At each iteration, N = 144 subsamples were taken, the same number of samples as in the original dataset.

To determine the relative timing of the movement of different body appendages, we fit Gaussian functions to the time derivative of the traces (Figure 1J-M, S1L-N). Derivatives were calculated using a Savitzky-Golay filter, with a window length of 0.16 s and a polynomial of order 1 (i.e., a straight line fit to the data in a sliding window of length 0.16 s). Gaussians were fit to data from 1 s before onset of the optomotor stimulus to 1 s into the stimulus period. Bounds were placed on the parameters such that *ω_p_* ∈ [0,100], *d* ∈ [0,1], and *t_p_* ∈ [0,1].

### METHODS DETAILS

#### Fly tethering

For optomotor response (Figure 1), silencing (Figure 3), unilateral activation (Figure 5), and bilateral activation experiments (Figure 6), female flies were anesthetized on a cold plate at 4°C and tethered to a tungsten wire (0.13 mm diameter) with UV-activated glue (Henkel; Bondic Inc.) at the anterior dorsal portion of the scutum. For bilateral activation experiments (Figure 6), we also fixed the head of each fly to its thorax by applying an additional drop of glue. Flies were allowed to recover for at least 15 minutes prior to testing.

For functional imaging experiments (Figure 4), we tethered each fly to a specially designed physiology stage that permitted access to the posterior side of the fly’s head (Maimon et al., 2008). We filled the holder with saline and removed a section of cuticle dorsal to the esophageal foramen, by first laser-cutting a window in the cuticle dorsal to the esophageal foramen using a custom built laser based on the design of (Huang et al., 2018; Valle et al.), then removing the window and dissected any remaining tissue obscuring the brain’s surface manually to improve imaging quality, including adipose bodies and trachea. This dissection protocol allowed us to simultaneously image neurons on the left and right side of the brain with limited damage to the animal.

#### Flight arenas and visual stimuli

For optomotor response (Figure 1, 3) and unilateral activation experiments (Figure 5), a tethered fly was positioned such that its stroke plane was horizontal and perpendicular to the vertical optical axis of a digital camera (Point Grey USB 3.0 camera equipped with a MLM3X-MP Computar macro lens and Hoya B-46RM72-GB IR-pass filter). An IR light source (driver: LEDD1B, and LED: M850L3; Thorlabs) illuminated the stroke planes of the left and right wing, and we used Kinefly (Suver et al., 2016) software to track the anterior-most angular excursion of the fly’s left and right wingbeats. To track the head, abdomen, and leg angles, we used DeepLabCut (Mathis et al., 2018). Training was based on the manual annotation of the joints of interest on two frames from each trial; the output of the trained network gave the x and y positions of each joint in pixels as a function of time. Flies were surrounded by a 11 x 3 panel (88 x 24 pixel) LED arena (470 nm) that covered 330° of azimuth with resolution of 3.75° in front of the fly (Reiser and Dickinson, 2008). Digital values for the wingbeat amplitudes were converted into voltages using a PhidgetAnalog 1002 and 1019 (Phidgets) and sent to a LED panel controller (IOrodeo) which was programmed so that flies could regulate the angular velocity of the visual display via the difference in wingbeat amplitude of the two wings for presentations of closed-loop stimuli, with a gain of 6° s^-1^ for each °ΔWBA. For presentations of closed-loop stripe fixation stimuli, a dark stripe subtending 15° was used. To induce the optomotor response, we rotated a striped drum (spatial frequency = 15°) in yaw at a velocity of 180° s^-1^. Three second presentations of yaw rotation were preceded and followed by 1 s of static striped drum to mitigate the effects of light artefacts and interspersed with 3 s of closed-loop stripe fixation. The direction of yaw rotation was shuffled pseudo-randomly. Each fly was subjected to up to 20 presentations of rotation in each direction.

Kinefly software was also used to track the wingbeat amplitude of the flies used for functional imaging experiments (Figure 4). We illuminated the wings using four horizontal fiber-optic IR light sources (M850F2, Thorlabs) distributed in a ~90° arc behind the fly. In these experiments, visual stimuli were presented using a 12 x 4 panel (96 x 32 pixel) arena that covered 216° of azimuth with a resolution of 2.25°, with a spectral peak of 450 nm to reduce light pollution from to LEDs into the photomultiplier tubes of the 2-photon microscope. We presented an array of visual starfield patterns, each for 3 s, alternated with 2 s static starfield patterns. Visual patterns were presented in pseudo-random order, including pitch, roll, and yaw in both directions and progressive and regressive motion.

For high-speed imaging experiments, flies were surrounded by a cylindrical array of 24 horizontal and 5 vertical 8×8 green LED panels with a red stage light filter (Roscolux Light Red no. 26) in front of the LEDs. Left and right wing shadows were recorded from the top view using a machine vision camera (Point Grey USB 3.0) recording at 30 Hz. The yaw position of a vertical black stripe (8 pixels wide) on a red background was controlled by the difference in left and right stroke amplitude, which was computed in real-time using Kinefly.

#### Anatomy

All anatomy panels were generated from 2-7 day old female *Drosophila melanogaster* unless otherwise specified. Full adult central nervous systems were dissected in 1xPBS, pH 7.4. All room temperature (RT) incubations are conducted with gentle agitation. Nervous systems were (1) fixed in 4% RT EM-grade paraformaldehyde for 25 minutes; (2) washed in RT 1XPBS 3X for 15 minutes and stored overnight in 4°C; (3) washed in RT 0.3% PBS-T (PBS containing 0.3% Triton-X) 5X for 15 minutes; (4) incubated in blocking buffer (7.5% normal goat serum in 0.3% PBS-T) for 1 hour at RT and transferred to 4°C overnight; (5) incubated in primary antibody, diluted in blocking buffer, at RT for 4 hours and transferred to 4°C for three nights; (6) washed 5X 15 min at RT in 0.3% PBS-T; (7) incubated with secondary antibody diluted in blocking buffer at RT for 4 hours and transferred to 4°C for an additional three nights; (8) washed 5X 15 min in RT 0.3% PBS-T; (9) mounted on a slide posterior side up with spacers using Vectashield Plus. For nervous systems with neuropil marker along with GFP and DsRED genetic reporters, we used for 1:30 nc82, 1:2000 chicken anti-GFP, and 1:2000 rabbit anti-DsRED primary antibodies. Secondary antibodies were then used at the following concentrations: anti-Mouse Cy5 (1:300), anti-Chicken Alexa 488 (1:1000), anti-Rabbit Alexa 546 (1:1000). Experiments with neuropil marker and Chrimson-Venus used the same antibody protocol without anti-DsRED primary and anti-Rabbit Alexa 546 secondary. Multicolor flip out experiments were conducted on 2 day old female flies; we used 1:30 nc82, 1:200 Rat anti-FLAG, and 1:300 Rabbit anti-HA primary antibodies, and 1:300 anti-Mouse Cy5, 1:500 anti-Rat Cy3, 1:500 anti-Rabbit Alexa 488 secondary antibodies.

Whole mount nervous systems were imaged using confocal microscopy [Zeiss LSM 880 using Zen Black (Carl Zeiss, Inc.)] Optical sections were imaged using a 40X water lens with 1-μM intervals, and 1024 × 1024 pixel resolution with tile scan. Post processing alignment of tiles was conducted using Zen Blue; anatomy panels were constructed in FIJI (Schindelin et al., 2012). In some panels, signal from off target single cells was digitally removed in FIJI to improve visualization of the neuron in question. Cell body quantification was conducted manually in FIJI. Cells were counted on each side of the brain and then averaged across 3 flies. If more than twice as many cells were counted on one side than the other, the brain was excluded from the count; this occurred twice, once each in the counts for SS02627 and SS01562.

#### Functional imaging

We imaged at an excitation wavelength of 930 nm using a gavanometric scan mirror-based two-photon microscope (Thorlabs) equipped with a Nikon CFI apochromatic, near-infrared objective water-immersion lens (40x mag., 0.8 N.A., 3.5 mm W.D.). We recorded tdTomato and GCaMP8m fluorescence in the posterior slope arbors of DNg02 neurons (bilaterally for data in Figure 4C-E; unilateral for data in Figure 4F-I). For bilateral imaging experiments, we acquired 72 x 29 μm images with 160 x 64 pixel resolution at 11.2 Hz, whereas we acquired 14.5 x 29 μm images with 64 x 128 pixel resolution at 13.1 Hz for unilateral imaging. To correct for motion in the x-y plane, we registered both channels for each frame by finding the peak of the cross correlation between each tdTomato image and the trial-averaged image. A region of interest (ROI) was defined by the 10% most variable GCaMP8m pixels. A centerline was defined manually and used to divide the ROI into left and right halves. Because the esophagus auto-fluoresces and appears in the GCaMP8m channel, we manually defined a mask to exclude it from the analysis. To correct for motion in z, we normalized the GCaMP8m fluorescence to the tdTomato fluorescence for a given frame by multiplying each pixel by the mean GCaMP fluorescence in the defined ROI for that frame and dividing by the mean tdTomato fluorescence in the ROI. For each side of the brain in each frame, we computed the fluorescence (F_t_) of the GCaMP8m signal by subtracting the average of the background from the average of the ROI. The background was defined as the mean fluorescence of the 5% dimmest pixels across the entire image. To standardize the measured neuronal activity across individual preparations, we normalized the baseline-subtracted fluorescence to the maximum observed for each individual fly on each side of the brain as ΔF/F = (F_t_ – F_0_)/F_95_, where F_0_ is the mean of the 5% lowest F_t_ and F_95_ is the mean of the 95% highest F_t_. For experiments in which we varied the rotational velocity of the visual display, we parsed the fluorescence data into bins of WBA, regardless of visual stimulus, by taking the mean WBA across 5 frames (Figure 4I). Data from a given fly was only included in the bin if there were at least 10 data points from that fly (i.e., if there were 10, 5 frame examples for the given fly at the WBA for a given bin). To be included in the dataset upon transition between quiescence and flight, both the quiescent period and the flight period were required to be longer than 1 s in duration, and the fly was required to have executed the transition at least 10 times.

#### Unilateral activation experiments

We positioned a fiber optic light guide (FT1500EMT; Thorlabs) behind the fly, aimed at the thorax, which conducted the output of a 617 nm LED (M617F1, Thorlabs) at ~3.4 mW/mm^2^. In each experiment, we elicited 10 responses to each stimulus duration with an interpulse interval of 10 s. For the data in Figure 5, stimulus durations of 0.05, 0.1, 0.3, 0.5, 1, 3, and 5 s were used. For the data in Figure 6, we used stimulus durations of 0.01, 0.025, 0.05, 0.1, 0.25, and 0.5 s. Durations were presented in pseudo-random order. The response to activation (Figure 5G-H) was determined as the average of the relevant quantity for the first 2 s following the onset of the stimulus minus the average for the second preceding the onset of the stimulus.

---

**Figure S1.**
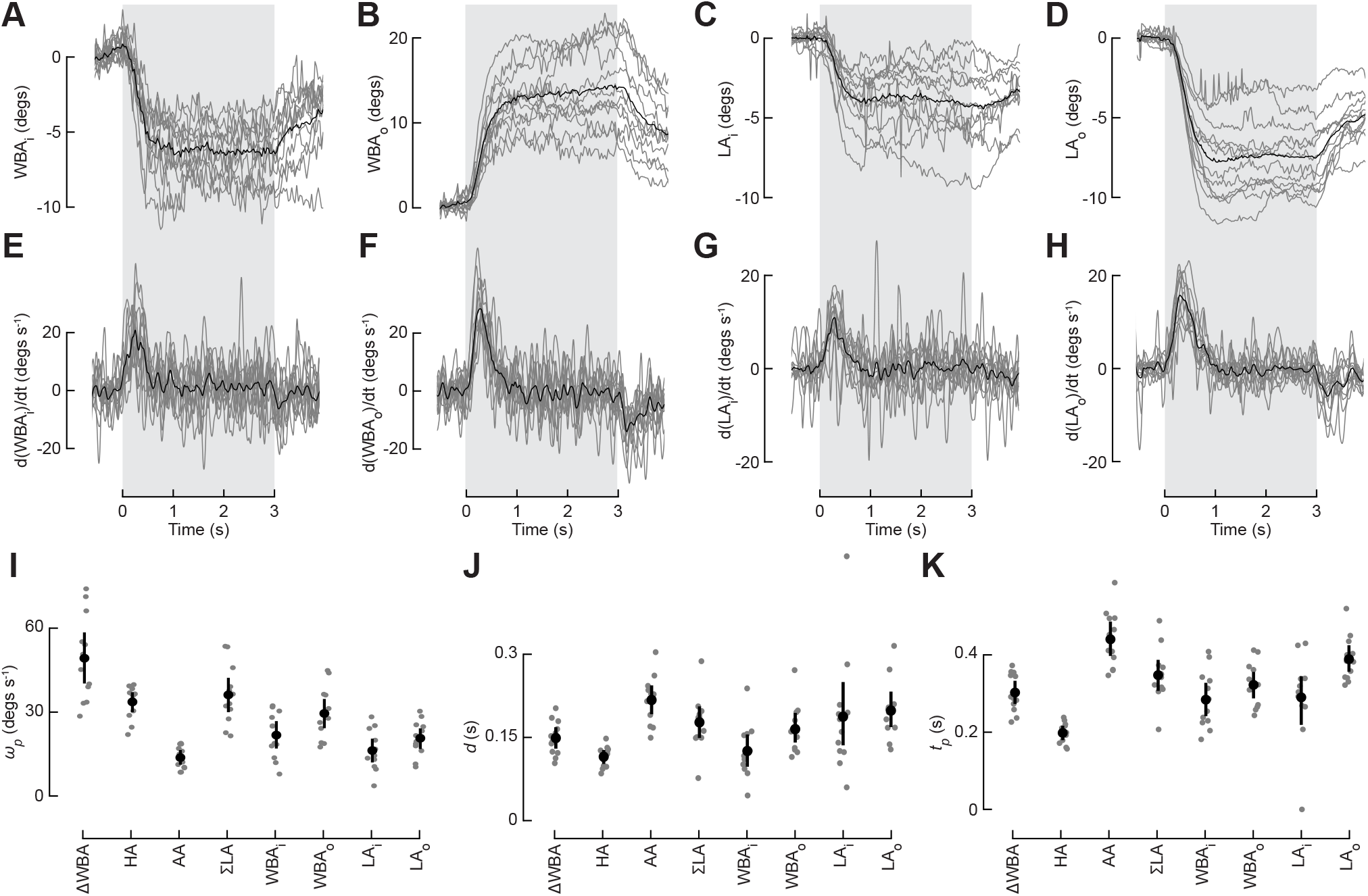
Wing and leg optomotor responses. (A) Averaged inside WBA (WBA_i_) during the presentation of yaw motion (n = 12 flies). Plotting conventions as in Figure 1D. (B) As in Figure S1A, for outside WBA, WBA_o_. (C) As in Figure S1A, for inside leg angle, LA_i_. (D) As in Figure S1A, for outside leg angle, LA_o_. (E) Time derivative of WBA_i_ during the presentation of yaw motion (n = 12 flies). Plotting conventions as in Figure 1D. (F) As in Figure S1E, for WBA_o_. (G) As in Figure S1E, for LA_i_. (H) As in Figure S1E, for LA_o_. (I) Peak angular velocity movements for all tracked kinematic parameters (ΔWBA, head, abdomen, ΣLA, individual wing, and individual leg angles). (J) As in Figure S1K, for duration of the movement. (K) As in Figure S1K, for the timing of the movement.

**Figure S2.**
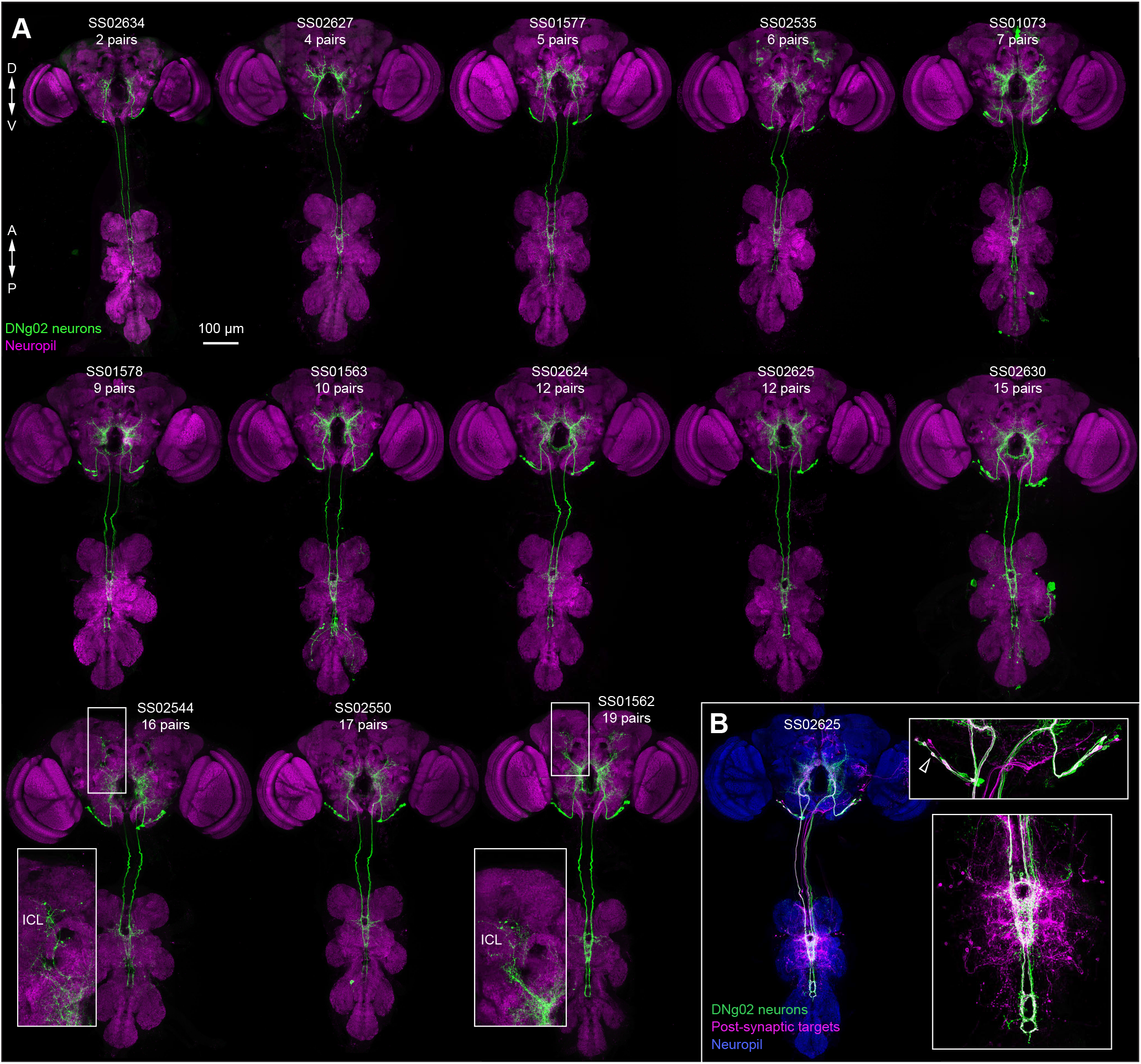
Full CNS morphology of DNg02 cells. (A-B) Confocal z-projections of DNg02 neurons in the fly central nervous system (CNS). Split-Gal4 ID numbers to visualize expression indicated in each panel. (A) Expression pattern in all driver lines targeting DNg02 neurons in the full CNS. Green: membrane GFP; magenta: nc82. Insets show projections to ICL present in a subset of the driver lines. (B) Postsynaptic targets of DNg02 neurons as revealed by trans-Tango in the full CNS. Green: membrane GFP in DNg02s; magenta: postsynaptic targets; blue: nc82. Top inset: DNg02 cell bodies. Coexpression of upstream and downsteam labeling suggesting reciprocal connectivity (empty arrowhead). Bottom inset: plexus of interneurons revealed in wing neuropil.

**Figure S3.**
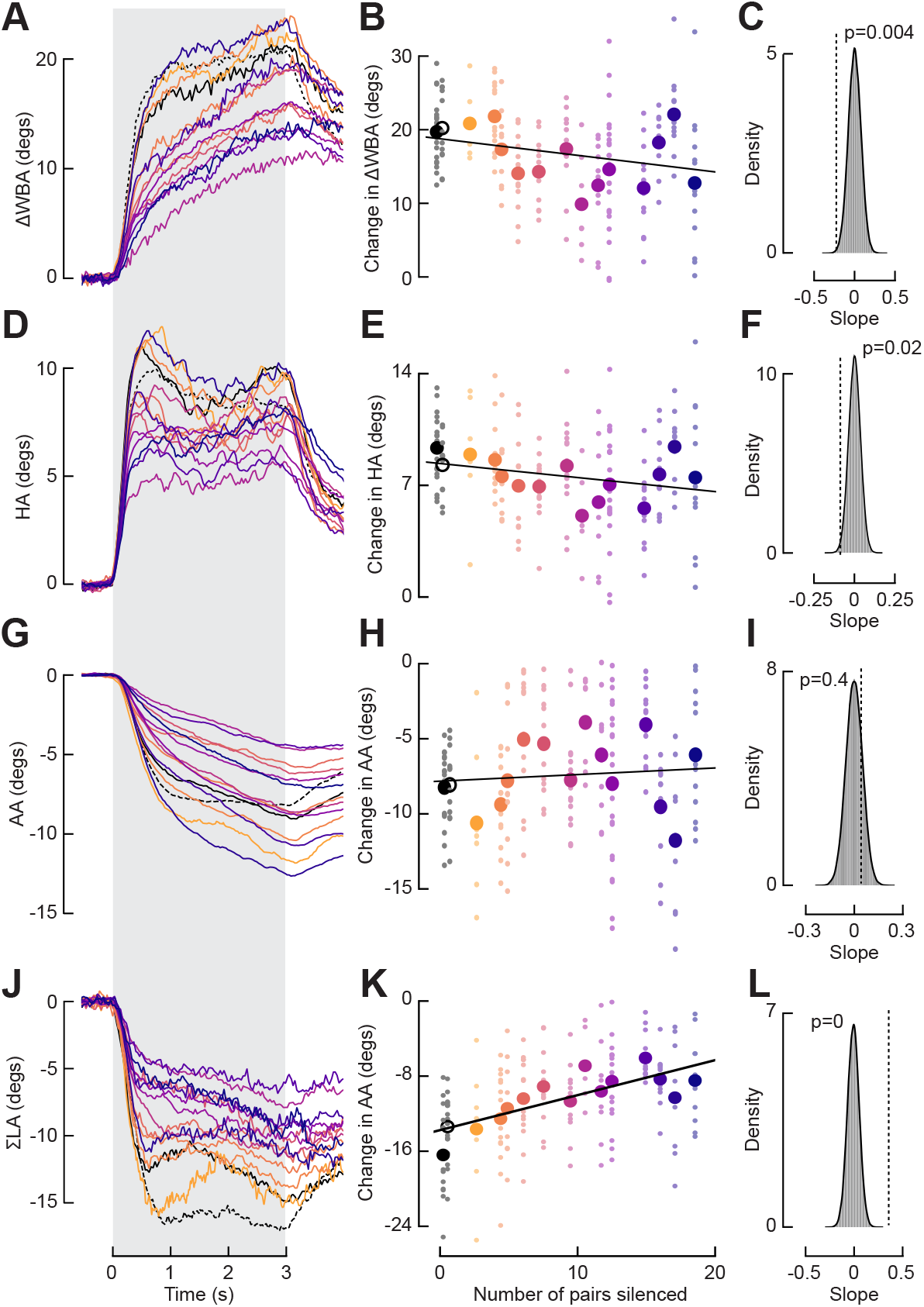
Results of silencing DNg02 neurons when lines with GNG projections are included. (A) Averaged ΔWBA during the presentation of yaw motion for flies with DNg02 neurons silenced with Kir (sample sizes throughout figure: empty-split vector control: n = 19; wild-type control: n = 12; SS02634: n = 7; SS02627: n =14; SS01577: n = 10; SS02535: n = 14; SS01578: n = 14; SS01563: n = 7; SS02625: n = 23; SS02624: n = 10; SS02630: n = 14; SS01073: n = 13; SS02550: n =17; SS02544: n = 12; SS01562: n = 13). Plotting conventions as in Figure 3A. Driver lines with GNG projections (SS01073, SS02550, SS02544, and SS01562) are included. (B) Change in ΔWBA plotted against the number of DNg02 pairs silenced. Plotting conventions as in Figure 3B (r^2^ = 0.17, based on mean values). (C) Distribution of 100,000 bootstrapped slopes of lines of best fit with responses and numbers of cells silenced randomly sampled with replacement for ΔWBA. Plotting conventions as in Figure 3C. (D) As in Figure S3A, for the head optomotor response. (E) As in Figure S3B, for the head optomotor response (r^2^ = 0.12, based on mean values). (F) As in Figure S3C, for the head optomotor response. (G) As in Figure S3A, for the abdomen optomotor response. (H) As in Figure S3B, for the abdomen optomotor response (r^2^ = 0.01, based on mean values). (I) As in Figure S3C, for the abdomen optomotor response. (J) As in Figure S3A, for the leg optomotor response. (K) As in Figure S3B, for the leg optomotor response (r^2^ = 0.63, based on mean values). (L) As in Figure S3C, for the leg optomotor response.

**Figure S4.**
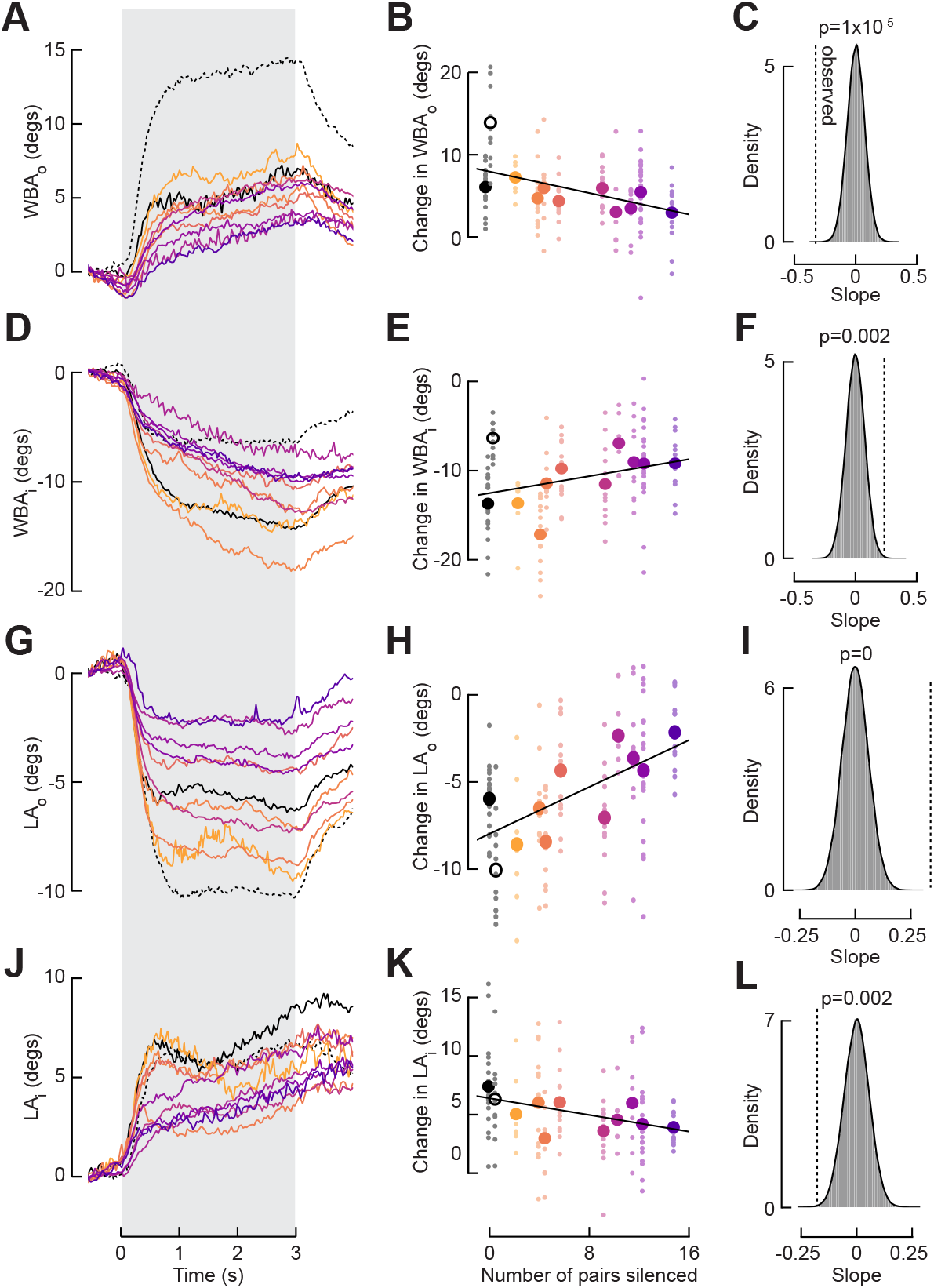
Silencing DNg02 neurons diminishes the magnitude of the optomotor response of individual wings and the legs. (A) Averaged WBA_o_ during the presentation of yaw motion for flies with DNg02 neurons silenced with Kir2.1 (sample sizes as in Figure 3). Plotting conventions as in Figure 3A. (B) Change in WBA_o_ plotted against the number of DNg02 pairs silenced. Plotting conventions as in Figure 3B (r^2^ = 0.43, based on mean values). (C) Distribution of 100,000 bootstrapped slopes of lines of best fit with responses and numbers of cells silenced randomly sampled with replacement for WBA_o_. Plotting conventions as in Figure 3C. (D) As in Figure S4A, for the WBA_i_ optomotor response. (E) As in Figure S4B, for the WBA_i_ optomotor response (r^2^ = 0.15, based on mean values). (F) As in Figure S4C, for the WBA_i_ optomotor response. (G) As in Figure S4A, for the LA_o_ optomotor response. (H) As in Figure S4B, for the LA_o_ optomotor response (r^2^ = 0.62, based on mean values). (I) As in Figure S4C, for the LA_o_ optomotor response. (J) As in Figure S4A, for the LA_i_ optomotor response. (K) As in Figure S4B, for the LA_i_ optomotor response (r^2^ = 0.30, based on mean values). (L) As in Figure S4C, for the LA_i_ optomotor response.

**Figure S5.**
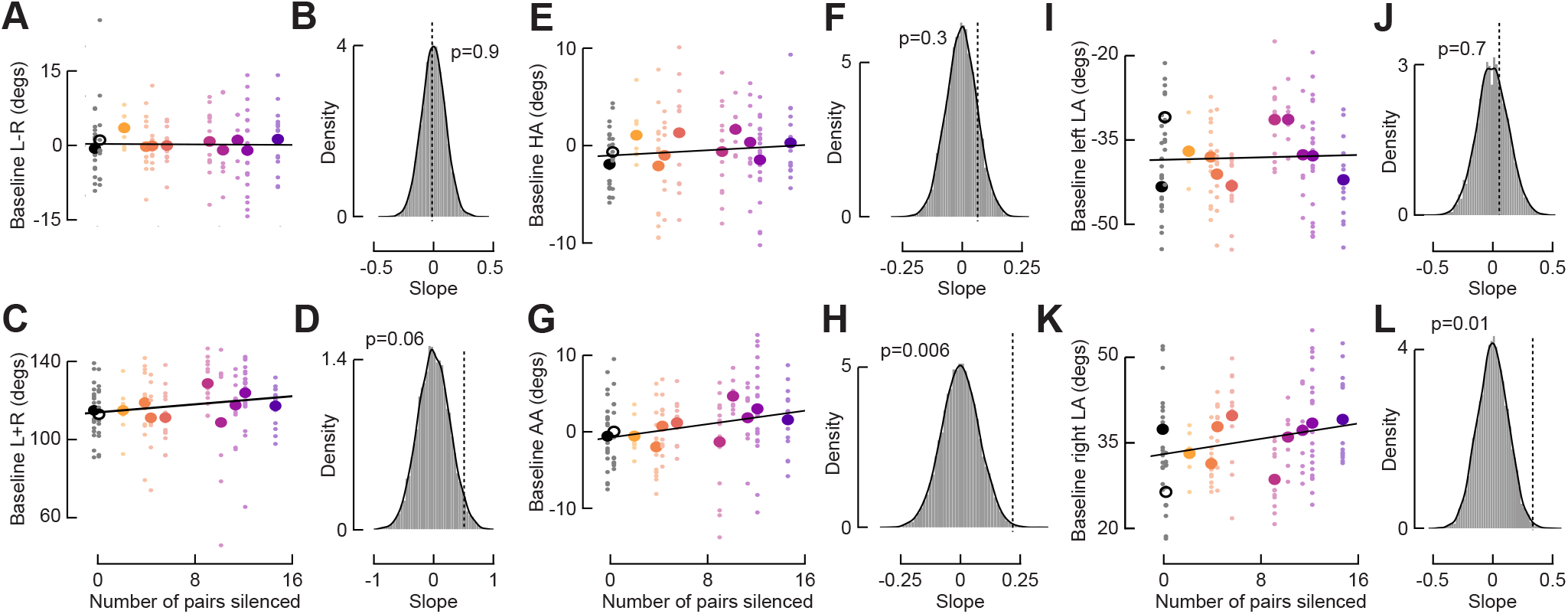
Effects of silencing DNg02 neurons on the optomotor response is not due to changes in baseline kinematics. (A) Baseline left minus right WBA plotted against the number of DNg02 pairs silenced (sample sizes as in Figure 3). The 1 s period of static grating preceding presentations of the optomotor stimuli is taken as the baseline period. Plotting conventions as in Figure 3B (r^2^ = 0.02, based on mean values). (B) Distribution of 100,000 bootstrapped slopes of lines of best fit with responses and numbers of cells silenced randomly sampled with replacement for baseline left minus right WBA. Plotting conventions as in Figure 3C. (C) As in Figure S5A, for WBA sum (r^2^ = 0.13, based on mean values). (D) As in Figure S5B, for WBA sum. (E) As in Figure S5A, for HA (r^2^ = 0.08, based on mean values). (F) As in Figure S5B, for HA. (G) As in Figure S5A, for AA (r^2^ = 0.34, based on mean values). (H) As in Figure S5B, for AA. (I) As in Figure S5A, for left LA (r^2^ = 0.001, based on mean values). (J) As in Figure S5B, for left LA. (K) As in Figure S5A, for right LA (r^2^ = 0.19, based on mean values). (L) As in Figure S5B, for right LA.

**Figure S6.**
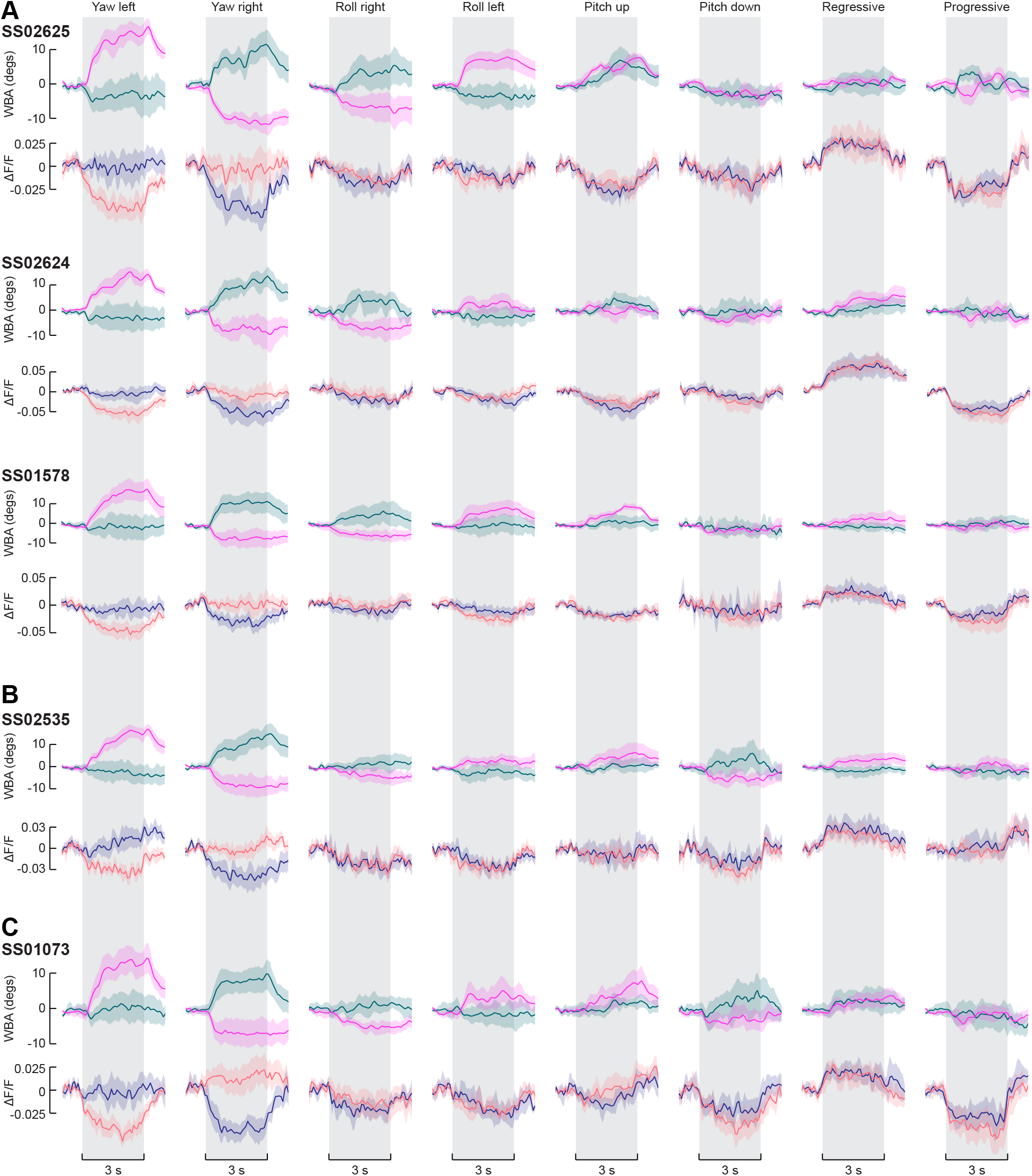
DNg02 activity is the same across different driver lines labeling different DNg02 variants. (A) Averaged wing and fluorescence responses to different patterns of optic flow for three driver line labeling both Type I and Type II DNg02 variants. Plotting conventions as in Figure 4D. (B) As in Figure S6A, for a driver line labeling exclusively Type II DNg02 neurons. (C) As in Figure S6A, for a driver line labeling Type II DNg02 neurons and some neurons with arborization in the GNG.

**Figure S7.**
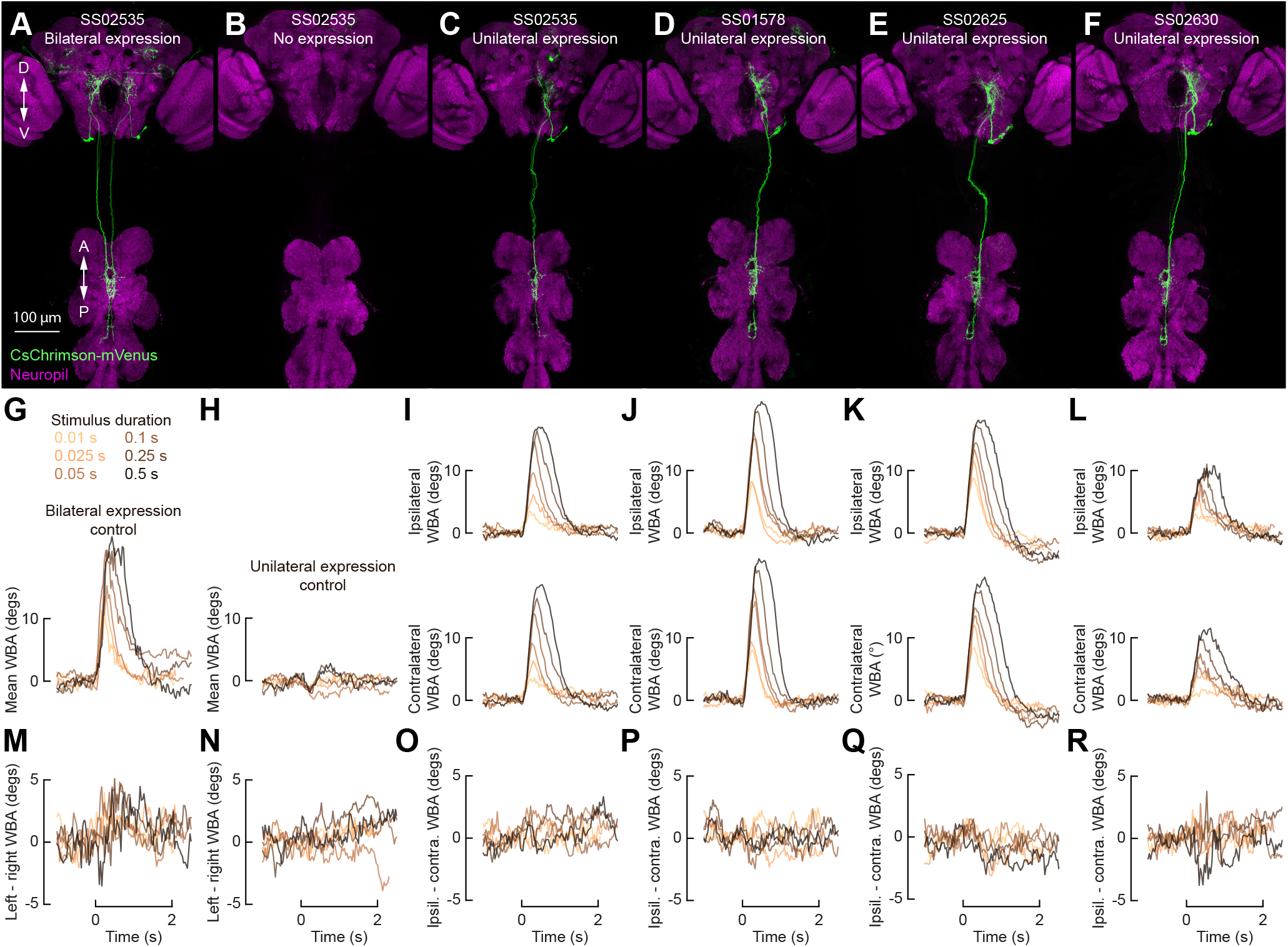
Unilateral DNg02 activation induces bilaterally symmetric wing responses at short stimulus durations and across driver lines. (A-F) Confocal z-projections of the fly central nervous system. Flp-recombinase under heat shock control coupled with 20XUAS-FRT>-dSTOP-FRT>-CsChrimson-mVenus enables temperature dependent, stochastic expression of CsChrimson-Venus (green) in DNg02 neurons bilaterally for SS02535 (A), on neither side for SS02535 (B), or unilaterally for SS02535 (C), SS01578 (D), SS02625 (E), and SS02630 (F). Magenta: nc82. (G) Baseline-subtracted mean WBA responses of flies (SS02535) with bilateral expression of CsChrimson to a red light stimulus of variable duration (0.01, 0.025, 0.05, 0.1, 0.25, or 0.5 s) (n = 6 flies). Flies were presented with a visual grating pattern in closed loop. (H) As in Figure S7G, for flies (SS02535) with no expression of CsChrimson (n = 16 flies). (I) Baseline-subtracted contralateral and ipsilateral WBA responses of flies (SS02535) with unilateral expression of CsChrimson (n = 15 flies). Plotting conventions as in Figure S7G. (J) As in Figure S7I, for SS01578 (n = 12 flies). (K) As in Figure S7I, for SS02625 (n = 14 flies). (L) As in Figure S7I, for SS02630 (n = 8 flies). (M) Mean WBA differential for the data in Figure S7G. Plotting conventions as in Figure S7G.(N) Mean WBA differential for the data in Figure S7H. (O) Ipsilateral minus contralateral differential for the data in Figure S7I.(P) Ipsilateral minus contralateral differential for the data in Figure S7J.(Q) Ipsilateral minus contralateral differential for the data in Figure S7K. (R) Ipsilateral minus contralateral differential for the data in Figure S7L.

